# Mesenchymal stem cell-extracellular vesicles deliver microRNAs that prevent nerve growth factor-induced sensory neuron sensitization

**DOI:** 10.64898/2026.01.12.698945

**Authors:** Lanhui Qiu, Paula-Milan Rois, Tuğdem Muslu-Ufuk, Jonathan L. Price, Alexander Cloake, Eric A. Miska, Tim L. Williams, Ewan St. John Smith

**Affiliations:** Department of Pharmacology, University of Cambridge; Department of Biochemistry, University of Cambridge; Department of Veterinary Medicine, University of Cambridge

## Abstract

Osteoarthritis (OA) affects 600 million individuals globally, pain being a hallmark symptom. Emerging clinical evidence supports the use of mesenchymal stem cells (MSCs) and their extracellular vesicles (MSC-EVs) for pain relief in knee OA. In mice, MSC-EVs ameliorate OA-induced pain and normalize knee-innervating neuron excitability. Moreover, it has been shown that overnight incubation of sensory neurons with MSC-EVs prevents the OA-associated mediator nerve growth factor (NGF) sensitizing sensory neurons. Here, we found that protease-mediated MSC-EV ‘shaving’ inhibited MSC-EV internalization into sensory neurons and the ability of MSC-EVs to prevent NGF-induced sensitization. In addition, acute, 10-minute, exposure of sensory neurons to MSC-EVs was also insufficient to counteract NGF. We hypothesized that MSC-EVs trigger transcriptional changes and found that inhibiting transcription prevented NGF-induced sensitization. MicroRNAs (miRNAs) can be delivered to cells by MSC-EVs, and certain miRNAs regulate transcription and pain; small RNA-sequencing of our MSC-EVs identified three candidate miRNAs, miR-21-5p, miR-148a-3p and miR-451a. Using gold nanoparticle delivery, each miRNA was able to prevent NGF sensitization of sensory neurons, a combination of all three showing the most pronounced effect. These findings demonstrate that MSC-EVs prevent NGF-induced sensory neuron sensitization via cellular uptake and transcriptional regulation that is mediated by miRNAs.

## Introduction

Osteoarthritis (OA) is a debilitating musculoskeletal disease affecting approximately 600 million people worldwide and is the 14^th^ leading cause for years lived with disability^1^. Joint pain is the hallmark symptom of OA, and inadequately managed pain limits joint function, reduces quality of life, and leads to long-term disability^2^. Current pharmacological treatments for OA pain, such as non-steroidal anti-inflammatory drugs and opioids, fail to provide sufficient relief and are often associated with adverse effects following chronic use^3^. Consequently, OA pain management remains a significant clinical challenge and underscores the need for disease-specific analgesics to address this unmet need.

Peripheral nociceptive input is a major contributor to OA pain. This is demonstrated clinically by reduced pain in OA patients following intra-articular administration of the local anesthetic lidocaine and by most, but not all, patients who undergo total knee replacement to remove the peripheral injury^4^. Preclinical studies reinforce this view: inhibition of nociceptive neuron activity with the quaternary anesthetic QX-314 ameliorates early OA pain^5^, and chemogenetic silencing of knee-innervating dorsal root ganglia (DRG) sensory neurons reverses inflammatory joint pain behaviors in mice^6,7^. Moreover, in the monosodium iodoacetate (MIA) and destabilization of the medial meniscus (DMM) OA models in mice, knee-innervating DRG sensory neurons become hyperexcitable, concomitant with pain behaviors^8,9^. In addition, in MIA treated rats, extracellular electrophysiological recordings reveal that knee-innervating sensory neurons become sensitized early after disease onset and remain so, whereas bone-innervating afferents become sensitized later in disease pathogenesis^10^. These findings highlight peripheral sensory neuron sensitization as a key process in OA pain and there is growing evidence for the plethora of cell-cell interactions and mediators that underpin this process^11,12^.

Among the molecular mediators implicated in OA pain, nerve growth factor (NGF) has emerged as particularly important. Not only does NGF cause sensitization when administered to many species^13,14^, but NGF is also present in the synovial fluid of humans with OA^15^, elevated in the knee joint in murine OA models^16^, and treatment with a soluble version of the NGF receptor, tropomyosin-related kinase receptor A (TrkA)^17^, anti-NGF antibodies^18^, or TrkA receptor inhibitors^19^ effectively suppresses pain-like behaviors in rodents undergoing models of OA. Although several anti-NGF antibodies have demonstrated clinical efficacy in OA pain management, concerns about their induction of rapidly progressive OA have prevented their regulatory approval^3^.

In parallel with the search for disease-modifying anti-OA drugs, mesenchymal stem cell (MSC) therapy has emerged as a promising disease modifying treatment. Clinical trials have reported pain relief and improved joint function in OA patients following MSC administration^20^. Although MSCs have shown encouraging results, enthusiasm for their clinical application is somewhat tempered by potential safety issues, such as the risk of tumor formation^21^. Consequently, extracellular vesicles released by MSCs (MSC-EVs) have been suggested as a safer alternative for OA therapy^22^, and a trial (NCT06431152) is currently investigating this in knee OA. It is thought that MSCs exert their effects primarily through paracrine mechanisms^23^, which makes MSC-EVs a key mediator of this paracrine action. Nano-sized MSC-EVs carry a complex cargo of proteins, lipids, and nucleic acids and offer a cell-free alternative to MSC therapy with fewer safety and logistical challenges^24^. Accumulating evidence shows that MSC-EVs reproduce many of the effects of the parent cell in OA models, including reduced inflammation, protection of cartilage, promotion of tissue repair, and, importantly, alleviation of pain^25–27^. Moreover, in the DMM mouse OA model, intra-articular injection of MSCs or MSC-EVs both caused significant reduction in pain-related behaviors without measurable improvements in joint histology^9^, indicating a likely direct action on nociceptive neuron activity rather than structural repair. Indeed, this hypothesis was confirmed when it was further demonstrated that knee-innervating neurons from MSC or MSC-EV treated DMM mice did not develop the hyperexcitability observed in saline treated DMM mice, and that the sensitizing effects of the OA-associated mediator NGF on cultured DRG neurons were prevented by treatment with MSC-EVs^9^.

Although MSC-EVs ameliorate DMM-induced pain and knee-innervating neuron hyperexcitability, and can normalize sensory neuron function in the presence of NGF^9^, the mechanisms underpinning these effects are unknown. In the present study, we investigated how MSC-EVs prevent NGF-induced DRG neuron sensitization. We observe that neuronal internalization of MSC-EVs is required for their full effect and that acute exposure of nociceptive neurons to MSC-EVs is insufficient to prevent NGF-induced sensitization, thus indicating that the effects of MSC-EVs might involve transcriptional changes, a hypothesis supported by experiments demonstrating that inhibiting transcription prevented the effect of MSC-EVs. Considering potential biomolecules that could be delivered to sensory neurons, small RNA-sequencing identified enriched microRNAs (miRNAs), some of which have been previously identified to regulate transcription and be involved in pain, including: miR-21-5p, miR-148a-3p, and miR-451a. Using gold nanoparticles to deliver miRNAs to sensory neurons, we find that a cocktail of all three recapitulates the ability of MSC-EVs to counteract the sensitizing effects of NGF, miR-21-5p being the most potent when test alone.

## Results

### 1. Establishing a high-throughput *in vitro* OA model for patch-clamp analysis using isolectin B4-ve DRG neurons

NGF is well-characterized as an agent that sensitizes sensory neurons^28,29^. Previous studies found that MSC-EVs act directly on DRG sensory neurons to prevent NGF-induced hyperexcitability, whereas exposing naïve DRG neurons to MSC-EVs had no significant impact on neuronal excitability^9^. NGF-induced sensitization involves binding to the TrkA receptor that is largely expressed by sensory neurons that do not bind the isolectin B4 (IB4)^30^, and we confirmed this TrkA vs. IB4 division by conducting immunohistochemistry on mouse lumbar L2-L5 DRG (those DRG that predominantly innervate the knee joint^7^); IB4 staining was observed in 43.8±4.8% of neurons, whereas TrkA was expressed by 36.7±3.5% of neurons, and, as previously observed^31,32^, there was a clear demarcation between the two neuronal populations, with minimal co-staining observed: only 8.5±1.8% of neurons co-stained for IB4 binding and TrkA expression (Fig. 1a). Therefore, to enhance the likelihood of neurons responding to NGF, we used live-cell imaging of IB4+ve neurons and made electrophysiological recordings from IB4-ve, i.e. likely TrkA expressing, neurons (Fig. 1b).

**Fig. 1:**
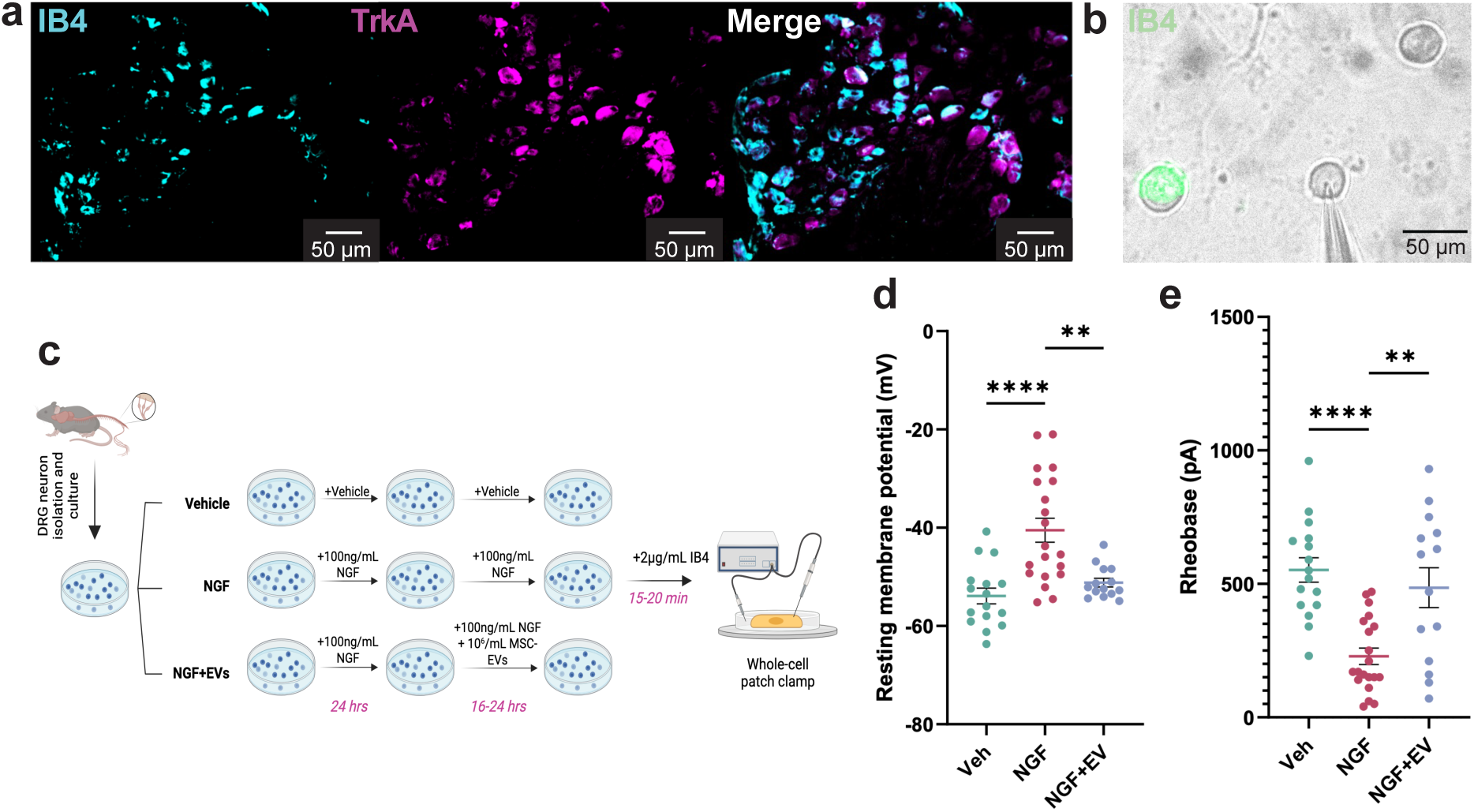
MSC-EV treatment prevents NGF-induced sensitization of IB4-ve DRG neurons. (**a**) Representative images of an L4 DRG section showing TrkA expression (magenta) and IB4 binding (cyan), and a merged image; DRG were from a 10-week-old male C57 mouse from which L2-L5 DRG sections were examined. IB4 staining was observed in 43.8±4.8% of neurons and TrkA expression in 36.7±3.5% of all neurons; data displayed as mean ± SEM. (**b**) IB4 staining is shown in green, recordings were made from IB4-ve neurons. (**c**) Schematic of experimental design: In vitro studies of DRG neurons performed in cultures with vehicle (control, n = 16 and N = 5; n = neurons and N = mice), 100 ng/ml NGF (n = 20 and N = 5), or NGF + MSC-EVs (10^6^/ml, n = 14 and N = 5), neurons being exposed to NGF for the whole culturing period in the second and third groups, MSC-EVs being added after 24-hours for the NGF + MSC-EVs group. DRG neurons were live-labelled with 2 µg/mL IB4-Alexa488 for 15-20-minutes at 37°C before patch clamp recordings. Only IB4-ve neurons were selected for electrophysiological studies. (**d**) Comparison of resting membrane potential (RMP) and (**e**) the minimum current required to induce an action potential (rheobase). Each dot represents an individual DRG neuron, horizontal lines with whiskers show the mean ± SEM. ** = P < 0.01 and **** = P < 0.0001 by one-way ANOVA followed by Tukey’s post hoc test.

Experimental groups were established as follows: (1) a control group with DRG neurons maintained in normal culture medium (water added as vehicle control), (2) an NGF group with DRG neurons maintained in culture medium with the addition of 100 ng/ml NGF to induce neuronal hyperexcitability as previously described^9^, and (3) an EV group in which DRG neurons were maintained in NGF-containing culture medium as in group 2, but with the addition of MSC-EVs after 24-hours, (Fig. 1c); MSC-EVs were prepared by differential ultracentrifugation, proteomic analysis confirming their identity (Supplementary Table 1). DRG neurons were labelled with 2 μg/mL IB4 for 15-20-minutes before making electrophysiology recordings of IB4-ve neurons. The capacitance of each neuron was recorded to ensure that similarly sized neurons were recorded from each group (Supplementary Table 2).

NGF-treated IB4-ve DRG neurons exhibited a significantly more depolarized resting membrane potential (RMP) compared to DRG neurons in the control group (control, -53.90±1.60 mV vs. NGF, - 40.51±2.45 mV, P<0.0001, Fig. 1d) and also exhibited a significantly lower rheobase (control, 551.88±45.94 pA vs. NGF, 228.50±30.87 pA, P<0.0001, Fig. 1e), thus, NGF sensitizes IB4-ve neurons. The RMP of DRG neurons in the EV group was significantly more negative than that of DRG neurons in the NGF group and was similar to that of DRG neurons in the control group (−51.18±0.87 mV in EV group, P<0.001 vs. NGF, Fig. 1d). Similarly, the rheobase of MSC-EV treated neurons was significantly higher than that of neurons in the NGF group (EV: 485.71±74.55 pA, P<0.001 vs. NGF, Fig. 1e). Therefore, administration of MSC-EVs to IB4-ve neurons prevents NGF-induced sensitization.

In NGF treated DRG neurons, action potentials were broader than those in the control group, also displaying a longer afterhyperpolarization duration (Supplementary Table 2). Importantly, as for RMP and rheobase, following MSC-EV treatment NGF no longer induced a significant change in either action potential half-peak duration or afterhyperpolarization duration (Supplementary Table 2). As RMP and rheobase represent the most robust and functionally relevant measures of intrinsic excitability in our preparation, these parameters were chosen as the primary readouts for subsequent analysis, further action potential parameter data being provided in supplementary tables.

These data therefore confirm that MSC-EVs prevent the sensitizing effects of NGF on IB4-ve DRG neurons enabling us to record from such neurons to determine the mechanism by which MSC-EVs produce their effects on sensory neurons.

### 2. Internalization of MSC-EVs by DRG neurons is required for robust prevention of NGF-induced sensitization

MSC-EVs can interact with other cells in multiple ways, for example, engaging in ligand-receptor interactions to trigger intracellular signaling or being internalized and delivering a variety of biomolecular cargo, from proteins to miRNAs^34^. We therefore firstly sought to examine the importance of extracellular surface receptor engagement internalization and/or intracellular cargo delivery in the effects of MSC-EVs.

Internalization of MSC-EVs can be blocked using inhibitors of endocytic pathways, for example, drugs that inhibit macropinocytosis (e.g. imipramine), clathrin- or dynamin-mediated endocytosis (e.g. chlorpromazine, dynasore, Pitstop 2), as well as those that disrupt lipid rafts or actin dynamics (e.g. methyl-β-cyclodextrin and cytochalasin D)^35^. However, most of these internalization-inhibiting compounds can themselves alter neuronal activity and thus are unsuitable for cleanly examining MSC-EV uptake in neurons when using an electrophysiological functional readout where ion channel activity is integral to what is being measured^35,36^. Consequently, we decided to perturb the ability of MSC-EVs themselves to be internalized, rather than inhibit processes in the recipient DRG neurons, using the proteases trypsin and proteinase K to cleave membrane proteins. Trypsin cleaves peptide bonds at the carboxyl side of lysine and arginine residues, and has been shown to selectively digest surface-accessible peptides while preserving MSC-EV membrane structure and functionality^37^. Proteinase K is a serine protease with minimal sequence specificity and efficiently degrades a wide range of surface proteins, including glycoproteins and membrane-associated proteins, without compromising MSC-EV morphology^38^. Treatment of MSC-EVs with either trypsin or proteinase K can therefore disrupt receptor-ligand interactions and impair internalization by recipient cells; while both enzymes effectively cleave surface proteins, their differing, but broad, substrate specificities means that they offer complementary approaches for probing MSC-EV internalization.

Firstly, we performed concentration and size distribution analyses of untreated MSC-EVs, trypsin-treated EVs (tEVs), and proteinase K-treated EVs (pkEVs) using nanoparticle tracking analysis (NTA, Fig. 2a): Untreated EVs exhibited a size distribution range of 80 to 520 nm with a modal size of approximately 152 nm, and a mean particle size of 176.8 nm, which aligns with previously reported findings for MSC-EVs^9,39^. In contrast, tEVs displayed a smaller size profile ranging from 50 to 420 nm, with two distinct peaks at ∼100 nm and ∼162 nm, and a reduced mean particle size of 122.8 nm, which aligns with the expected effect of trypsin-mediated surface protein cleavage. The emergence of a second peak may suggest differential susceptibility among EV subpopulations, with some vesicles retaining protease-resistant surface proteins that may preserve receptor-ligand interactions and internalization capacity. Proteinase K treatment had a similar impact on EV morphology, such that pkEVs showed a size distribution with a range of 20 to 320 nm, a modal size of 129 nm and a mean particle size of 142.3 nm (Supplementary Table 3).

**Fig. 2:**
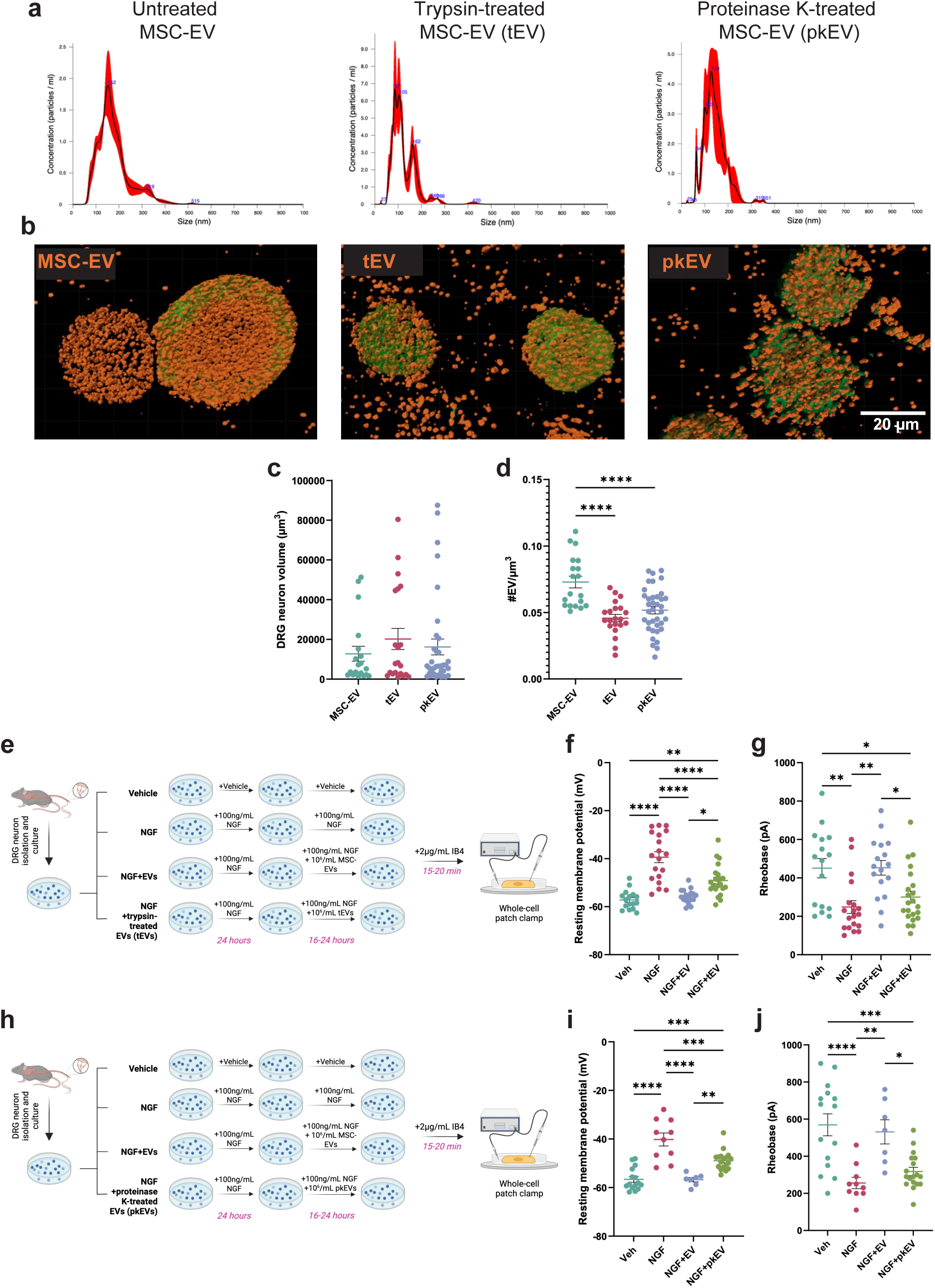
Internalization of MSC-EVs by DRG neurons is required for prevention of NGF-induced sensitization. (**a**) Particle distributions of untreated MSC-EVs, trypsin-treated EVs (tEVs), and proteinase K-treated EVs (pkEVs). (**b**) Confocal images showing reduced internalization of tEV and pkEV by DRG neurons (indicated by red, PKH26 puncta in upper images) with greater fluorescent signal observed in the extracellular space rather than within the neuronal soma; tEV or pkEV were co-cultured with DRG neurons for 10-minutes at 37 °C prior to imaging. (**c**) Distribution of DRG neuron soma volume and (**d**) quantification of internalized MSC-EVs per DRG neuron µm^3^, i.e. EV density, following 10-minute incubation; each dot represents an individual DRG neuron. (**e**) Schematic of experimental design for tEVs. In vitro studies of DRG neurons performed in cultures with vehicle (control, n = 16 and N = 7, n = neurons and N = mice), 100 ng/ml NGF (n = 19 and N = 7), NGF + MSC-EVs (10^6^/ml, n = 17 and N = 7), NGF + tEVs (10^6^/ml, n = 22 and N = 7). Neurons were exposed to NGF for the whole culturing period in the second, third and fourth groups, MSC-EVs and tEVs were added after 24-hours for the third and fourth groups respectively. (**f**-**g**) Comparison of action potential parameters: (**f**) RMP and (**g**) rheobase. (**h**) Schematic of experimental design. In vitro studies of DRG neurons performed in cultures with vehicle (control, n = 15 and N = 5, n = neurons and N = mice), 100 ng/ml NGF (n = 10 and N = 5), NGF + MSC-EVs (106 /ml, n = 7 and N = 5) or NGF + pkEVs (10^6^ /ml, n = 18 and N = 7). Neurons were exposed to NGF for the whole culturing period in the second, third and fourth groups, MSC-EVs and pkEVs were added after 24-hours for the third and fourth groups respectively. For all electrophysiology recordings, DRG neurons were live-labelled via incubation with 2 µg/mL IB4-Alexa488 for 15-20-minutes at 37°C and only IB4-ve neurons were selected for electrophysiological studies. Horizontal lines with whiskers show the mean ± SEM. * = P < 0.05, ** = P < 0.01, *** = P < 0.001 and **** = P < 0.0001 by one-way ANOVA followed by Tukey’s post hoc test.

High-resolution images of MSC-EVs were acquired by transmission electron microscopy (TEM) and demonstrated that MSC-EVs had the spherical shape and characteristic lipid bilayer structures consistent with their known morphology of EVs^9,26^, i.e. shaving had no significant impact on EV morphology and lipid bilayer integrity (Supplementary Figure 1) consistent with previous reports^40,41^.

We next used confocal microscopy to visualize the internalization of MSC-EVs by DRG neurons, for both non-treated EVs and for those that have been ‘shaved’ with trypsin or proteinase K. To do this, MSC-EVs were labelled with PKH26, a red fluorescent membrane dye that enables efficient tracking, visualization, and quantification of vesicles within the cellular environment^42,43^. PKH26-labelled MSC-EVs were applied to dissociated DRG neurons for 10 minutes before imaging (Fig. 2b). To quantify uptake, we measured the number of internalized EVs within individual DRG neurons using a machine learning pipeline, which provides robust, quantitative data on EV internalization. No significant differences were observed in the volumes of DRG neurons that were analyzed (Fig. 2c), but, consistent with example confocal images (Fig. 2b), the density of internalized EVs was significantly reduced in both shaved EV groups compared to the control group (control: 0.073±0.004 EVs/μm^3^; tEV: 0.046±0.003EVs/μm^3^, P<0.0001 vs. control; pkEV: 0.052±0.003 EVs/μm^3^, P<0.0001 vs. control) (Fig. 2d).

To determine if less efficient internalization impacted the ability of MSC-EVs to prevent NGF-induced sensitization, we conducted two sets of experiments as outlined in Fig. 1, but with the addition of a fourth group in each set, either tEV or pkEV (i.e. neurons treated with NGF+tEVs or NGF+pkEVs, Fig. 2e-j); on each day of experiments, recordings were made from neurons in multiple conditions to prevent bias resulting from how neurons from a particular mouse responded to treatments. The capacitance of neurons was recorded, and average values were not different between groups in either the tEV or pkEV experiment (Supplementary Tables 4 and 5).

NGF+tEV-treated DRG neurons had a significantly more depolarized RMP compared to NGF+EV-treated neurons (EV, -55.67±0.71 mV vs. tEV, -49.12±1.36 mV; P < 0.0001) and control neurons (−57.18±0.99 mV; P<0.01), but the RMP was, however, still significantly more hyperpolarized compared to NGF alone (NGF, -39.38±2.92 mV vs. tEV, -49.12±1.36 mV, Fig. 2f, Supplementary Table 4), thus, although impacting neuronal activity, tEVs are not as effective as untreated EVs in counteracting the effects of NGF on RMP. Neurons from the tEV group also exhibited a significantly lower rheobase compared to neurons from the EV group and control group, which was not significantly different to purely NGF-treated neurons (EV, 452.84±38.05 pA, vs. tEV, 300.00±30.72 pA, P<0.05; vs. control, 450.63±50.27 pA, P<0.05; NGF, 248.42±32.98 pA, Fig. 2g, Supplementary Table 4). Thus, as with the RMP results, tEVs cannot fully ameliorate the sensitizing effects of NGF. As was observed with tEVs, NGF+pkEV-treated DRG neurons had a significantly more depolarized RMP than NGF+EV-treated neurons (EV, -56.75±0.90 mV vs. pkEV, -48.90±0.93 mV, P<0.01) and control neurons (−56.60±1.18 mV, P<0.001), but the RMP was significantly more hyperpolarized compared to that of NGF-treated neurons (−40.20±2.65 mV, P<0.001, Fig. 2I, Supplementary Table 5); thus, like tEVs, pkEVs are not as effective as untreated EVs in preventing NGF-induced RMP depolarisation. In addition, NGF+pkEV-treated DRG neurons exhibited a significantly lower rheobase than NGF+EV-treated neurons and control neurons, which was not significantly different to that of purely NGF-treated neurons (EV, 531.43±65.08 pA vs. pkEV, 317.78±22.56 pA P<0.05; vs. control 569.33±59.20 pA; P<0.001; NGF, 256.00±30.56 pA; Fig. 2j, Supplementary Table 5). Therefore, as was observed with tEVs, pkEVs, both of which display less efficient DRG neuron internalization (Fig. 2d), are unable to prevent NGF-induced DRG neuron sensitization.

### 3. Transcriptional regulation is required for MSC-EVs to counteract NGF-induced DRG neuron sensitization

Given that overnight incubation with MSC-EVs prevented NGF-induced sensitization in IB4-ve DRG neurons, but that internalization happens within 10-minutes (Fig. 2b,d), we next examined whether a 10-minute incubation would be sufficient to counteract the sensitizing effects of NGF. This experiment enabled us to test the hypothesis of whether the effects of EVs observed were more likely to result from rapid post-translational modification (e.g. due to delivery of a second messenger or engagement of plasma membrane receptors to induce intracellular signaling), or comparatively slower transcription.

Three groups of neurons were analyzed: control, NGF-treated and NGF+10-minute MSC-EV treatment (Fig. 3a). However, unlike overnight EV incubation (Fig. 1c-e), 10-minute MSC-EV incubation did not significantly affect either the RMP or rheobase of DRG neurons compared to those in the NGF group, both the RMP and rheobase being significantly different to those of control neurons (Fig. 3b,c, Supplementary Table 6). These results suggest that rapid cell signaling events, such as post-translational modification are unlikely to play a major role in how MSC-EVs modulate sensory neuron excitability. This finding is consistent with clinical observations, where patients receiving MSC-EV treatment typically experience pain relief and functional improvement over longer time frames^44,45^.

**Fig. 3:**
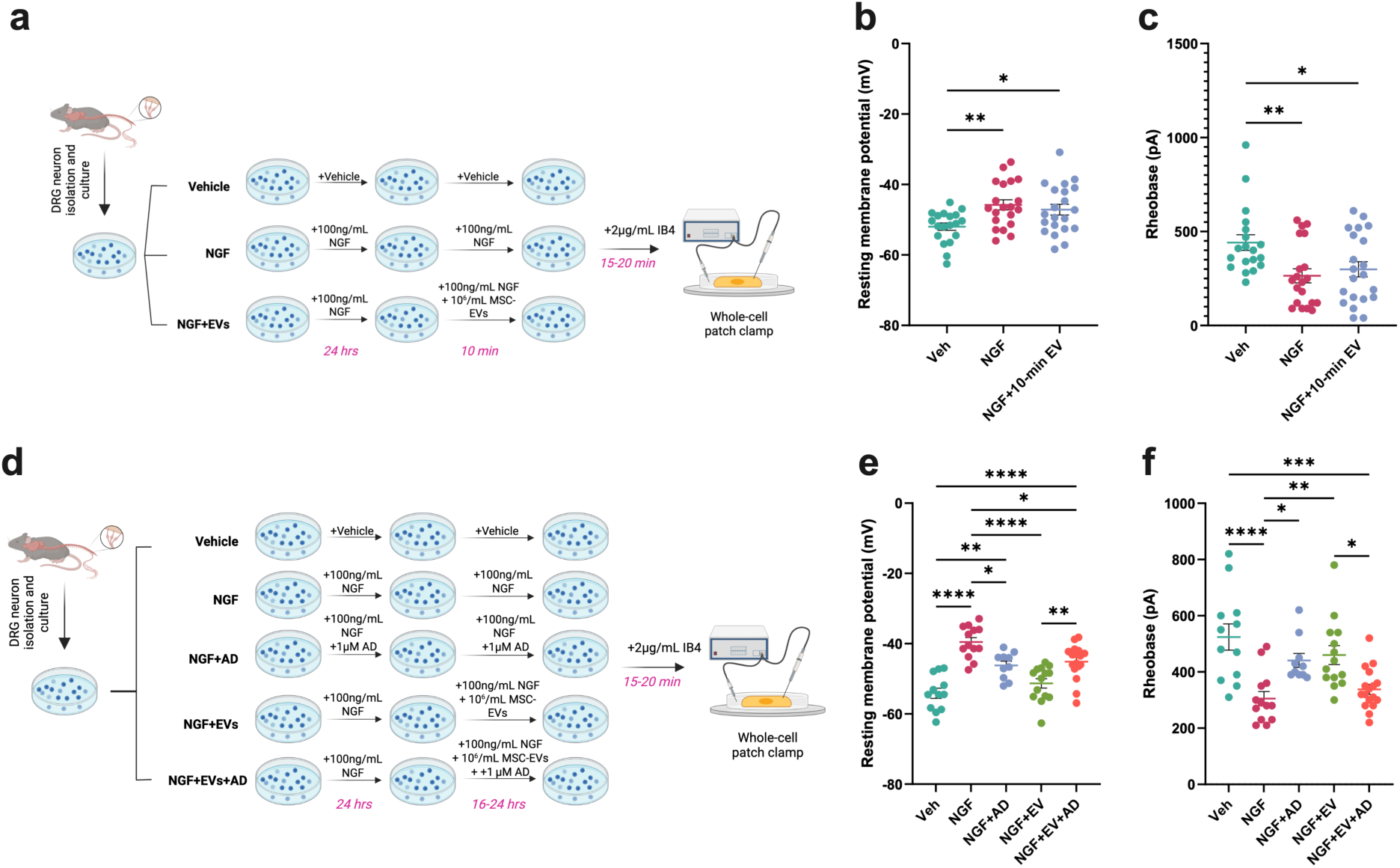
Transcriptional regulation is required for MSC-EVs to fully counteract NGF-induced neuronal sensitization. (**a**) Schematic of experimental design for assessing the effects of a 10-minute MSC-EV treatment on NGF-induced sensitization. *In vitro* studies of DRG neurons were performed in cultures with vehicle (control, n = 20 and N = 5, n = neurons and N = mice), 100 ng/ml NGF (n = 19 and N = 5) or NGF + MSC-EVs (10^6^/ml, n = 21 and N = 5) for 10-minutes. (**b**-**c**) Comparison of action potential parameters: (**b**) RMP and (**c**) rheobase. (**d**) Schematic of experimental design assessing the impact of AD on the ability of MSC-EVs to reverse NGF-induced sensitization in IB4-negative DRG neurons. *In vitro* studies of DRG neurons performed in cultures with vehicle (control, n = 12 and N = 7, n = neurons and N = mice), 100 ng/ml NGF (n = 13 and N = 7), NGF+AD (n = 10 and N = 7), NGF + MSC-EVs (10^6^/ml, n = 14 and N = 7), or NGF+MSC-EVs+AD (NGF+EV+AD, n = 17 and N = 7). (**e**-**f**) Comparison of action potential parameters: (**e**) RMP and (**f**) rheobase. For all electrophysiology recordings, DRG neurons were exposed to NGF for the whole culturing period in groups 2-5, AD being present in group 2 throughout the whole culturing period, EVs or EVs+AD being added to neurons for the groups 4 and 5, respectively. DRG neurons were live-labelled via incubation with 2 µg/mL IB4-Alexa488 for 15-20 min at 37°C and only IB4-ve neurons were selected for electrophysiological studies. Each dot represents an individual DRG neuron sample. Horizontal lines with whiskers show the mean ± SEM. * = P < 0.05, ** = P < 0.01, *** = P < 0.001 and **** = P < 0.0001 by one-way ANOVA followed by Tukey’s post hoc test.

To directly determine if MSC-EVs might exert their effects through transcriptional regulation, we applied the transcriptional inhibitor actinomycin D (1 μM, AD) to DRG neuronal cultures. NGF regulates neuronal excitability in part via transcriptional mechanisms^46^, therefore, co-treatment with AD was anticipated to interfere with NGF-induced DRG neuron sensitization, as well as any potential impact on EV function. We thus analyzed five groups of DRG neurons (Fig. 3d): (1) control, (2) NGF-treated, (3) NGF + AD, (4) NGF + MSC-EVs, and (5) NGF + MSC-EVs + AD (AD applied only during the second overnight incubation). Neurons were exposed to NGF for the whole culturing period in groups 2-5, neurons in group 3 were also exposed to actinomycin D for the whole culturing period. For groups 4 and 5, actinomycin D and/or MSC-EVs were added after 24 hours. We hypothesized that if MSC-EVs primarily exert their effects through transcriptional regulation, then treatment with AD would abolish their ability to reverse the effects of NGF, i.e., the NGF+MSC-EV group would show reduced excitability compared to NGF alone, but this effect would not occur in the NGF+MSC-EV+AD group.

Consistent with previous results, compared to control neurons, NGF-treated DRG neurons exhibited a significantly more depolarized RMP (control: -54.13±1.46mV vs. NGF: -39.53±1.26 mV, P<0.0001, Fig. 3e, Supplementary Table 7) and a significantly lower rheobase (control: 524.17±46.60pA vs. NGF: 305.38±25.13pA, P<0.0001, Fig. 3f). As expected, the RMP and rheobase of the NGF + AD group (RMP: -46.20±1.20 mV; Rheobase: 441.00±24.88 pA) were significantly different from those of the NGF-only group (P<0.05); however, they also remained significantly different from control neurons with regard to RMP (P<0.01), i.e. NGF still induced sensitization in the presence of AD, but not to the same extent as when AD was absent, which likely reflects NGF’s modulation of neuronal excitability through both transcription-dependent and transcription independent mechanisms^47^. Both RMP and rheobase in the NGF+EVs+AD group were significantly different from the NGF+EVs group (RMP: NGF+EVs+AD: -45.15±1.20 mV vs. NGF+EVs: -51.31±1.34 mV, P<0.01; Rheobase: NGF+EVs+AD: 338.24±17.75 pA vs. NGF+EVs: 460.00±33.55 pA, P<0.05, Fig. 3e-f). Whilst AD prevented MSC-EVs from fully normalizing rheobase, the value for NGF treated neurons not being significantly different to that of those in the NGF+EVs+AD group, the RMP in the NGF+EV+AD group remained significantly different from NGF alone (−39.53±1.26 mV, P<0.05 vs. EV+AD, Fig. 3e). This suggests that although transcriptional regulation is necessary for MSC-EVs to exert their full effects, part of their action may also involve transcription-independent mechanisms.

### 4. A cocktail of miR-21-5p, miR148a-3p and miR451a fully reverses NGF-induced sensory neuron sensitization

Results described above suggest that the effects of MSC-EVs on neuronal excitability are not immediate, but rather likely involve a slower process, such as transcription. Given the diverse biomolecular cargo of MSC-EVs and their known capacity to modulate transcriptional activity, we focused our investigation on their microRNA (miRNA) content. miRNAs are small, highly conserved non-coding RNA molecules that can regulate gene expression, several miRNAs having been shown to reduce pain and transcription^48–51^.

To determine the specific contribution of candidate miRNAs in MSC-EVs to DRG neuron modulation, we first performed small RNA sequencing to determine the miRNA profile of our MSC-EVs (Fig. 4a). Three miRNAs were selected for further investigation based upon their high enrichment in MSC-EVs (miR-21-5p, miR-148a-3p, and miR-451a; 3^rd^, 2^nd^, and 13^th^ most highly abundant, respectively) and their known involvement in pain modulation and transcriptional regulation^51–56^. One challenge was to optimize a technique for delivering miRNA to DRG neurons because mouse DRG neurons are highly susceptible to being killed by standardly used lipid-based transfection reagents. Indeed, initial trials with Lipofectamine RNAiMAX, which can be used to deliver miRNA, siRNA etc. to cells, demonstrated that this reagent was toxic to DRG neurons (data not shown), and thus an alternative method was required. We tested a gold nanoparticle delivery system, which has been used to deliver miRNAs to muscle stem cells^57^. To assess the effects of individual miRNAs on DRG neuron excitability, each candidate miRNA was conjugated to gold nanoparticles. Four types of nanoparticles were synthesized, firstly, Cy5-labelled gold nanoparticles (Cy5NPs) where the particles are functionalized with a non-coding Cy5-labelled oligonucleotide, enabling fluorescent tracking of nanoparticle uptake. Unlike the miRNA-loaded nanoparticles, Cy5NPs do not carry functional miRNA cargo; however, the comparable oligonucleotide length and surface chemistry enable them to mimic the physical and structural aspects of miRNA delivery, albeit without biological activity. Thus, Cy5NPs serve as a control to assess internalization efficiency and nanoparticle behavior independently of miR-mediated effects^58^. The other three types synthesized were: miR-21-5p-conjugated nanoparticles (miR-21-5pNPs), miR-148a-3p-conjugated nanoparticles (miR-148a-3pNPs), and miR-451a-conjugated nanoparticles (miR-451aNPs).

**Fig. 4:**
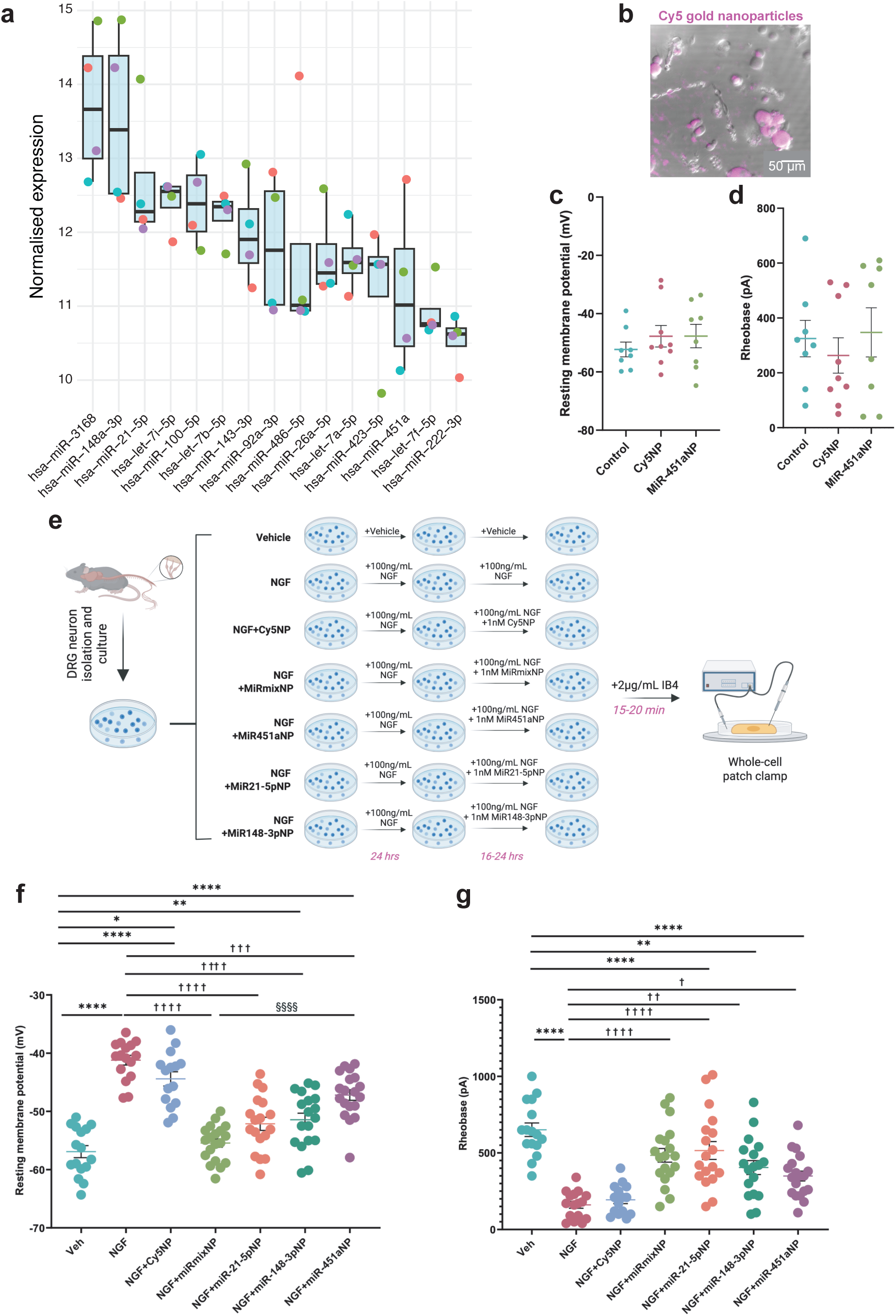
An miRNA cocktail of miR-21-5p, miR148a-3p and miR451a prevents NGF-induced DRG neuron sensitization. (**a**) miRNA profiling of MSC-EVs (n = 4), each dot represents a different sample. **(b)** Oligonucleotide-Cy5-conjugated gold nanoparticles (Cy5NP) are incorporated into DRG neurons with very high efficiency. (**c**-**d**) Neither Cy5NPs nor miR-451a alter the resting membrane potential or action potential parameters of DRG neurons. In vitro studies of DRG neurons were performed in cultures with vehicle (control, n = 8 and N = 2, n = neurons and N = mice), 1nM Cy5NP (n = 9 and N = 2) or 1nM miR451aNP gold nanoparticle (n = 8 and N = 2). Comparison of action potential parameters: **(c)** RMP and (**d**) rheobase. (**e**) Schematic of experimental design for miRNA-conjugated gold nanoparticles. In vitro studies of DRG neurons were performed in cultures with vehicle (control, n = 16 and N = 6, n = neurons and N = mice), 100 ng/ml NGF (n = 16 and N = 6), NGF + 1nM Cy5NP (n = 15 and N = 6), NGF + 1nM miRmixNP (n = 19 and N = 6), NGF + 1nM miR451aNP (n = 19 and N = 6), NGF + 1nM miR21-5pNP (n = 18 and N = 6) or NGF + 1nM miR148-3pNP (n = 18 and N = 6). In NGF treated groups, DRG neurons were exposed to NGF for the whole culturing period Cy5NP or miRNAs being added to neurons as indicated after 24-hours. DRG neurons were live-labelled via incubation with 2 µg/mL IB4-Alexa488 for 15-20-minutes at 37°C and only IB4-ve neurons were selected for electrophysiological studies. Total number of neurons recorded in each group is summarized in the table. (**f**) Comparison of RMP and (**g**) rheobase. Each dot represents an individual DRG neuron recording. Horizontal lines with whiskers show the mean±SEM. ^*/†^ = P < 0.05, ^**/††^ = P < 0.01, ^***/†††^ = P < 0.001 and ^****/††††/§§§§^ = P < 0.0001 by one-way ANOVA followed by Tukey’s post hoc test.

We first confirmed that gold nanoparticles underwent internalization into DRG neurons using Cy5NPs and confocal microscopy, overnight incubation with 1 nM Cy5NPs demonstrating highly efficient nanoparticle internalization in DRG neurons (Fig. 4b). We then assessed, as a proof of principle, whether nanoparticles themselves influenced neuronal excitability. For these initial experiments, miR-451aNPs were chosen at random and compared alongside Cy5NPs and untreated DRG neurons. No significant differences were observed in action potential parameters across the three groups (Fig. 4c-d, Supplementary Table 8), indicating that neither nanoparticles nor miR-451aNPs significantly altered basal neuronal activity. Thus, gold nanoparticles can be used to efficiently deliver miRNA to DRG neurons, and this delivery method appears to have no overt impact on neuronal excitability.

Next, we combined the three miRNA-conjugated gold nanoparticles into a single miRNA cocktail to assess their collective effect on NGF-induced DRG neuron sensitization, as well as simultaneously testing the effects of each individual miRNA (Fig. 4e). The total concentration of the miRNA cocktail was kept equivalent to that used for each individual miRNA treatment, ensuring direct comparability between groups. Consistent with previous data, NGF-treated DRG neurons exhibited a significantly depolarized RMP compared with controls (Control: -56.91 ± 1.04 mV vs. NGF: -41.21 ± 0.83 mV, P < 0.0001, Fig. 4f) and a markedly reduced rheobase (Control: 651.25 ± 44.17 pA vs. NGF: 160.63 ± 22.37 pA, P < 0.0001, Fig. 4g). Cy5NP treatment did not reverse the effects of NGF (Cy5NP: RMP: -44.41 ± 1.22 mV; Rheobase: 194.00 ± 25.09 pA), consistent with previous findings showing that gold nanoparticles alone do not alter sensory neuron excitability (Figure 4c-d).

Unlike Cy5NP and similarly to EVs, the miRNA cocktail fully reversed the effects of NGF on DRG neurons, restoring both RMP (−55.44 ± 0.72 mV, *P* < 0.0001 vs. NGF, Fig. 4f) and rheobase (484.21 ± 44.68 pA, *P* < 0.0001 vs. NGF, Fig. 4G) to values not significantly different from control neurons. Each individual miRNA also attenuated NGF’s effects, although to varying degrees. Regarding RMP, miR-21-5pNP (−52.13 ± 1.13 mV, *P* < 0.0001 vs. NGF), miR-148a-3pNP (−51.43 ± 1.12 mV, *P* < 0.0001 vs. NGF), and miR-451aNP (−47.19 ± 0.91 mV, *P* < 0.001 vs. NGF) all produced significant recovery compared with NGF alone (Fig. 4f). Similarly, rheobase was increased relative to NGF following treatment with miR-21-5pNP (515.56 ± 58.23 pA, *P* < 0.0001 vs. NGF), miR-148a-3pNP (404.44 ± 45.63 pA, *P* < 0.01 vs. NGF), and miR-451aNP (348.95 ± 32.34 pA, *P* < 0.05 vs. NGF) (Fig. 4g). Among the individual treatments, miR-21-5p was the most effective, yielding RMP and rheobase values statistically indistinguishable from both the miRNA cocktail and control neurons (Fig. 4f-g). By contrast, miR-451a produced the weakest effect, with the RMP remaining significantly more depolarized compared with the miRNA cocktail (Fig. 4f-g). miR-148a-3p showed an intermediate effect, improving both parameters relative to NGF, but not achieving the same efficacy as miR-21-5p (Fig. 4f-g, Supplementary Table 9).

## Discussion

OA is a poorly treated condition, with no reliable treatment for modifying disease pathogenesis or for effectively managing the pain that individuals living with OA experience. Managing OA pain is critical considering the impact that living with chronic pain has on everyday life^59–61^. Previous work has shown that MSC-EVs reduce knee-innervating neuronal hyperexcitability and pain behaviors in a mouse OA model, as well as preventing NGF-induced sensitization of cultured DRG neurons^9^. These findings align with what has been observed from studies where humans living with OA have been given MSCs^62^. The overarching aim of this study was to determine the mechanisms by which MSC-EVs counteract the sensitizing effects of NGF and thus identify potential therapeutic avenues. We find that sensory neuron internalization of MSC-EVs and subsequent miRNA delivery are key to the ability of MSC-EVs to prevent NGF-induced sensitization.

Despite species differences, IB4+ve and IB4-ve neurons represent two broadly distinct subpopulations of sensory neurons with notable differences in anatomy, function, molecular profile^32,63^, as well as electrophysiological evidence highlighting functional divergence between these subsets^31^. Although previous studies have shown that MSCs and MSC-EVs reduce sensory neuron hyperexcitability and pain behaviors in murine OA^9^, the mechanisms by which this occurs remain unknown. By selectively recording from IB4-ve DRG neurons, we enhanced the probability that recordings would be made from neurons likely responsive to NGF because as our data confirmed what others have documented previously, i.e. that in mouse DRG TrkA (i.e. NGF responsive neurons) positive neurons are largely IB4-ve^30,31,64^. In this study, we firstly confirmed our previous observation that incubation with NGF sensitizes DRG neurons, i.e. compared to control IB4-ve DRG neurons, NGF-treated IB4-ve DRG neurons had a more depolarized RMP and lower rheobase, and that these effects can be prevented by overnight incubation with MSC-EVs^9^.

The internalization of MSC-EVs is essential for many of their functional impacts on recipient cells^65,66^. Studies have shown that MSC-EVs can attenuate fibrosis and promote tissue repair primarily through the uptake and delivery of their molecular cargo^66^. In contrast, naïve MSCs that do not internalize EVs fail to exhibit these differentiation effects, highlighting the critical role of EV internalization in driving cellular changes^67^. This is because once target cells take up MSC-EVs, their cargo, including proteins, nucleic acids, and lipids, can interact with intracellular components and signaling pathways, leading to functional changes in the recipient cells^68^. We therefore sought to determine if EV internalization plays a role in EV prevention of NGF-induced sensitization of DRG neurons. We observed that a 10-minute incubation is sufficient for MSC-EVs to be internalized by DRG neurons, but that internalization is significantly less efficient when MSC-EVs are treated with either trypsin or proteinase K to remove extravesicular and surface proteins. Moreover, such ‘shaved’ MSC-EVs were unable to fully replicate the effects of untreated MSC-EVs. For example, the NGF-induced decrease in DRG neuron rheobase is reversed by EVs, but this reversal is not observed following incubation with either tEVs or pkEVs. Moreover, although the NGF-induced depolarisation of DRG neuron RMP was partially reversed by tEV or pkEV treatment, the RMP of these neurons was still significantly more depolarized compared to control neurons, whereas untreated EVs fully prevent the NGF-induced RMP depolarisation. These findings suggest that biomolecules within MSC-EVs, which function intracellularly, play a crucial role in mediating their effects on sensory neuron function, and/or that EV-receptor signaling is important in mediating the reversal of NGF-induced sensitization.

An EV-delivered biomolecule could initiate rapid intracellular signaling, such as post-translational modification of ion channels underpinning action potential generation, or induce slower cellular changes, such as modifying transcription of such ion channels. When exposing DRG neurons to an acute, 10-minute EV incubation, there was no prevention of NGF-induced sensitization, which suggests that neither rapid EV receptor-ligand signaling nor delivery of a fast-signaling intracellular messenger plays a major role on how EVs counteract NGF-induced sensitization of DRG neurons. By contrast, the fact that overnight incubation of DRG neurons with EVs is required for their NGF counteracting effects suggests that transcription might be involved. To support this hypothesis, we observed that inhibition of transcription strongly reduced the effects of MSC-EVs. Taken together, these results suggest that the transfer of intracellular cargo is likely essential for mediating the observed effects on DRG neurons.

In recent years, miRNAs contained in MSC-EVs have attracted considerable interest due to their ability to modulate pain by regulating the expression of target genes involved in nociceptive signaling pathways. For example, in neuropathic pain, pro-inflammatory macrophages release EVs that are enriched in miR-155 and these EVs are taken up by sensory neurons in the DRG, where miR-155 binds to the 3′UTR of SHIP1 mRNA, leading to post-transcriptional suppression of SHIP1 expression. This reduction in SHIP1 protein decreases its phosphatase activity, resulting in enhanced PI3K/Akt signalling and increased interleukin-6 expression, which contributes to mechanical hypersensitivity^69^. Similarly, EVs have been reported to carry miR-let-7b, which activates TLR7-TRPA1 signalling in neurons and glial cells. This interaction promotes Ca^2+^ influx and increases excitatory synaptic transmission within nociceptive pathways, thereby contributing to pain hypersensitivity^67^. By contrast, miR-30b alleviates oxaliplatin-induced neuropathic pain by direct post-transcriptional suppression of the voltage-gated sodium channel Nav1.6 (SCN8A) in DRG neurons, thereby reducing Na^+^ influx, suppressing neuronal hyperexcitability, and ultimately diminishing pain hypersensitivity^70^.

Profiling of MSC-EVs in this study identified high levels of several miRNAs that have been associated with both pain modulation and transcriptional regulation, including miR-21-5p, miR-148a-3p, and miR-451a. We first established that oligonucleotides can be delivered to DRG neurons in vitro via conjugation to gold nanoparticles, thus demonstrating an efficient way to deliver miRNAs to cells that are well-known to be susceptible to lipid-based oligonucleotide delivery methods that work well in most cells. We observed that overnight incubation of a cocktail of all 3 miRNAs, i.e. as would be delivered via MSC-EVs, was able to recapitulate the effects of EVs themselves, and that miR-21-5p was most potent when each miRNA was administered individually. At the molecular level, miR-21-5p can directly bind to the 3′UTR of PDCD4 mRNA, leading to its post-transcriptional downregulation. This reduction in PDCD4 expression removes its inhibitory effect on anti-inflammatory IL10 gene transcription, thereby promoting anti-inflammatory cytokine production^71^. Importantly, in vivo studies in rats demonstrate that administration of miR-21-5p mimics can alleviate neuropathic pain behaviors, with evidence that this occurs through direct post-transcriptional downregulation of pro-inflammatory mediators^56^. Taken together, these findings support the hypothesis put forward by our data that miR-21-5p-enriched MSC-EVs could contribute to the observed reduction in DRG neuron excitability by engaging transcriptional regulatory pathways to counteract NGF. miR-148a-3p is highly expressed in rat DRG neurons and regulates transcriptional programs relevant to neuronal survival and plasticity^72^. However, miR-148a-3p has also been implicated in pain, much of the published work emphasizing its role in microglial modulation, for example, FOXA2/miR-148a-3p/SMURF2 signaling in neuropathic pain^51^. In addition, studies in central nervous system neurons have shown neuron-intrinsic regulation of miR-148a-3p during neuroinflammation^73^ and miR-148a-3p is predicted to bind to the 3′UTR of DNMT1 mRNA, leading to post-transcriptional downregulation of DNMT1, a key transcriptional regulator implicated in spinal mechanisms of neuropathic pain^74^, suggesting that it can influence sensory-neuron transcriptional plasticity. miR-451a has been less extensively studied in pain, but recent evidence demonstrates a direct neuronal role. In a pancreatic cancer pain model, tumor-derived EVs suppressed DRG miR-451a expression, leading to upregulation of the neuropeptide VGF and increased neuronal excitability^75^. Since VGF is a well-established mediator of pain sensitization, this identifies a plausible miR-451a-VGF axis within DRG neurons, whereby EV-delivered miR-451a could suppress VGF activity. Additionally, miR-451a has been implicated in regulating transcriptional stress responses and neuronal survival pathways^76^, supporting a broader role in neuronal resilience. Taken together, these findings substantiate that MSC-EV cargo miRNAs have the potential to modulate pain not only through indirect immune and inflammatory mechanisms, but also via sensory neurons themselves.

In conclusion, this study demonstrates that MSC-EVs can suppress the sensitizing effects of NGF on DRG neurons, with their efficacy dependent on internalization and transcriptional regulation. Importantly, we identify EV-enriched miRNAs: miR-21-5p, miR-148a-3p, and miR-451a, that can modulate sensory neuron excitability thereby offering mechanistic insight into how MSC-EVs achieve their analgesic effects in vivo, as well as opening the door to future studies examining the potential for miRNA-mediated pain management in OA.

## Authorship Statement

The study was conceived and designed by Lanhui Qiu, Prof. Tim Williams, and Prof. Ewan St. John Smith. Dr Paula Milan-Rois conducted MSC-EV RNA isolation and AuNP miRNA functionalization. Dr Jonathan Price assisted with the analysis of RNA-seq results. All other data were collected and analyzed by Lanhui Qiu. The paper was drafted by Lanhui Qiu and Prof. Ewan St. John Smith with all authors assisting with editing and approving the final version.

## Acknowledgements

This work was supported by the David James Trust Fund (LQ), Wellcome Trust (EStJS, 225856/Z/22/Z), a joint and equal investment from UKRI and Arthritis UK (EStJS, MR/W002426/1), and Arthritis UK (AC and EStJS, RG21973). We thank Dr Luke Pattison for assistance with confocal imaging, Dr Robin Antrobus at the Cambridge Institute for Medical Research Proteomics Facility (CIPF) for conducting LC-MS/MS analysis, Dr Karin Mueller and colleagues at the Cambridge Advanced Imaging Centre for assistance with electron microscopy, and Dr James Boyd and Darran Clements for assistance with Airyscan microscopy.

## Materials and Methods

### Animals

All animal experiments were performed in accordance with the Animals (Scientific Procedures) Act 1986 Amendment Regulations 2012 under Project Licenses granted to E.St.J.S. (P7EBFC1B1 and PP5814995) by the Home Office and approved by the University of Cambridge Animal Welfare Ethical Review Body. Experiments were performed using a mixture of male and female C57BL6/J mice (10-15 weeks old) purchased from Envigo and housed conventionally with nesting material and a red plastic shelter in a temperature-controlled room at 21 °C, with a 12-hour light/dark cycle and access to food and water ad libitum. Animals were humanely killed by CO_2_ exposure followed by cervical dislocation.

### Dorsal root ganglia neuron isolation and culture

Dorsal root ganglia (DRG) neuron isolation and culture were performed as previously described^77^. In brief, before collecting DRG, poly-D-lysine glass-bottomed dishes (MatTek, USA) were coated with laminin (1mg/mL, Invitrogen UK, 23017-015) and incubated at 37°C for 1-hour before removing excess laminin, washing with H_2_O and drying. Those lumbar DRG that predominantly innervate the knee (L2-L5) were collected post-mortem and placed into cold dissociation medium (L-15 Medium (Gibco™ UK, 31415-086) with 1.5% NaHCO_3_ (Gibco™ UK, 25080-094)). DRG were enzymatically digested in prewarmed collagenase solution (dissociation medium with 1 mg/mL collagenase (Sigma-Aldrich UK, C9891) and 6 mg/mL bovine serum albumin (BSA, Sigma-Aldrich UK, A2153)) for 15 mins followed by digestion in trypsin solution (dissociation medium with 1 mg/mL trypsin (Sigma-Aldrich UK, T9935), 6 mg/mL BSA, Sigma, UK) for 30 mins at 37°C before mechanical trituration with a P1000 tip 20-25 times in culture medium (dissociation medium with 1% penicillin/streptomycin (Gibco UK, 15140-122) and 10% fetal bovine serum (Sigma-Aldrich UK, F7524). Following trituration, brief centrifugation (1000 g, 30s) was used to separate remaining DRG tissue from dissociated neurons. The resulting supernatant, containing the dissociated neurons, was collected in a separate tube, and then 2 mL of culture medium was added to pelleted DRG neurons for further trituration. Trituration and brief centrifugation were repeated 5 times until 10 mL of supernatant was collected. Collected supernatant was centrifuged at 1000 g for 5 mins and the pellet was resuspended in culture medium and 100 L plated into the center of each glass-bottomed dish. Neurons were incubated at 37°C with 5% CO_2_ for 2 hours before then adding 1.9 mL of culture medium.

Neurons were cultured in either standard medium (control) or medium supplemented with recombinant mouse NGF-β (100 ng/mL, SRP4304, Sigma-Aldrich) to model OA-associated sensitization *in vitro*. The β isoform represents the mature, biologically active subunit of NGF that binds TrkA and initiates downstream signaling and was therefore used for experimental induction of sensitization^78^. After 24 h, cultures were refreshed with medium containing either vehicle (water, negative control), NGF (100 ng/mL, positive control), or NGF in combination with additional treatments as specified in individual figure legends. Neurons were then maintained for the required duration prior to electrophysiological recordings.

In all EV-related experiments, a concentration of 10^6^ EVs/mL was used. Actinomycin D (AD; BS-0481A, BioServUK) was applied at a final concentration of 1 µM in relevant experiments. In experiments involving miRNA-conjugated gold nanoparticles, a final concentration of 1 nM was used. Where possible, the experimenter was blinded to the culturing conditions.

### Immunohistochemistry

Immunohistochemistry was performed as previously described^79^. Briefly, L2-L5 DRG were collected and post-fixed for 1h in 4% paraformaldehyde (PFA, pH 7.4), followed by overnight incubation in 30% (w/v) sucrose at 4 °C for cryoprotection. DRG were next embedded in Shandon M-1 Embedding Matrix (Thermo Fisher Scientific), snap frozen in 2-methylbutane (Honeywell International) on dry ice and stored at -80 °C. Embedded DRG were sectioned (12 μm) using a Leica Cryostat (CM3000; Nussloch, Germany), mounted on Superfrost Plus microscope slides (Thermo Fisher Scientific) and stored at - 20°C until staining.

Slides were defrosted, washed with phosphate buffered saline (PBS)-Tween and blocked in antibody diluent solution: 0.2% (v/v) Triton X-100, 5% (v/v) donkey serum and 1% (v/v) bovine serum albumin in PBS for 1-hour at room temperature before overnight incubation at 4°C with anti-TrkA (goat polyclonal, 1:1000; R&D Systems, AF1056) primary antibody. Following incubation, slides were washed 3×10 minutes with PBS-T and then exposed to anti-goat Alexa Fluor 568 (1:200 Jackson ImmunoResearch) secondary antibodies for 2-hours at room temperature. Isolectin B4 conjugated to Alexa Fluor 488 (IB4-AF488, Invitrogen I21411) was applied at a 1:100 dilution for 15 minutes at 37 °C for immunohistochemistry. Slides were washed in PBS-Tween, mounted and imaged with an Olympus BX51 microscope (Tokyo, Japan) and QImaging camera (Surrey, Canada). Exposure times were kept constant during image acquisition.

Images were analyzed using ImageJ. Brightness and contrast adjustments were applied uniformly across all images. Negative controls without the primary antibody showed no detectable staining. An automatic “minimum error” threshold algorithm was applied to 8-bit images to distinguish background from objects. Background subtraction was performed by measuring the average signal intensity in negative control areas and subtracting it from the experimental regions of interest. The mean grey value of each neuron in a DRG section was measured and normalized between the highest and lowest intensity neuron for that section^80^. The threshold used for scoring a neuron as positive for a stain was set as the normalized minimum grey value across all sections +2 times standard deviation (SD). IHC results are reported as mean ± standard error of the mean (SEM)

### Patch clamp electrophysiology

DRG neurons were first incubated for 15-20-minutes at 37°C with 0.5 μg/mL isolectin B4 (IB4) conjugated to either Alexa Fluor 488 (Invitrogen I21411) or Alexa Fluor 568 (Invitrogen I21412). Excess fluorophore was removed by washing the neurons twice with ECS prior to beginning patch-clamp recordings. Glass patch pipettes (3-6 MΩ, Hilgenberg) were pulled using a P-97 Flaming/Brown puller (Sutter Instruments, USA) from borosilicate glass capillaries and loaded with intracellular solution (ICS), which (in mM) contained KCl (110), NaCl (10), MgCl_2_ (1), EGTA (1), HEPES (10), Na_2_ATP (2), Na_2_GTP (0.5) adjusted to pH 7.4 with KOH and osmolality adjusted to 300-310 mOsm using sucrose. DRG neurons were bathed in extracellular solution (ECS) (in mM): NaCl (140), KCl (4), CaCl_2_ (2), MgCl_2_ (1), glucose (4), HEPES (10), adjusted to pH 7.4 with NaOH, and osmolality was adjusted to 280-295 mOsm using sucrose. Cells were observed using an Olympus IX70 Fluorescence Microscope. Solutions were applied with a gravity-driven multi-barrel perfusion system (Automate Scientific). Data was acquired with a Multiclamp 700A amplifier (Molecular Devices, USA) and digitized via Digidata 1440A digitizer (Molecular Devices, USA).

For current-clamp recordings, bridge-balance compensation was used to reduce steady-state voltage errors. Data were sampled at 20kHz and filtered at 5kHz. Step current (−100 pA to 2000 pA) for 80 ms in 10 pA steps was injected to generate action potentials (APs) under current-clamp mode. All recordings were made using a Multiclamp 700A amplifier in combination with Clampex (Molecular Devices, USA) software. Pipette and membrane capacitance were compensated, with series resistance being compensated by ∼70% to minimize voltage errors. IB4+ve neurons were visualized by observation of Alexa-488 or Alexa-568 fluorescence (CellCam Kikker 100MT) and recordings made from IB-ve neurons.

Data were assessed for normality using the Shapiro-Wilk test, and parametric or non-parametric statistics were used as appropriate. Patch clamp results were analyzed with one-way ANOVA with Tukey’s post-hoc test. Statistical analysis and graph generation were carried out in GraphPad Prism 8.0 software (USA), Biorender®, R studio® and Python. Statistical significance was defined as *p* ≤ 0.05.

### Extracellular vesicle isolation and characterization

#### Cell lines and culture

Mesenchymal stem cells (MSC, Lonza, Cat # PT-2501) were cultured and MSC extracellular vesicles (MSC-EVs) collected and enriched as previously described^9^. Passage four human bone marrow-derived MSCs were cultured in α-MEM culture medium (Thermo Fisher Scientific, UK, 12571063) supplemented with 10% v/v fetal calf serum (Thermo Fisher Scientific, UK), 1% (v/v) glutamax (100×) (Gibco, UK), 1% (v/v) penicillin/streptomycin (Gibco, UK), and incubated at 37 °C, 5% CO_2_. After 3 days from seeding into T175 flasks, reaching approximately 80% confluence, cells were switched to serum-free culture medium (α-MEM (Thermo, UK), 1% (v/v) glutamax (100×) (Gibco, UK), 1% (v/v) penicillin/streptomycin (Gibco, UK)) for 48-hour incubation.

#### EV isolation: ultracentrifugation (UC) and size-exclusion chromatography (UF-SEC)

Extracellular vesicles were isolated from the MSCs by combined ultracentrifugation (UC) and size-exclusion chromatography (UC-SEC) method^9,81^. Conditioned medium, defined as cell culture medium that has been incubated with live cells and thus contains the secreted EVs of those cells, was collected after 48-hours of incubation with MSCs in serum-free medium to allow for EV release. Before EV isolation, the medium was clarified by centrifugation at 300 × g for 5-minutes to remove cells and large debris. The supernatant was removed and further centrifuged at 2,000 g for 20-minutes, 4°C. Supernatant was then transferred into polycarbonate ultracentrifuge tubes (Beckman, USA) for differential sequential UC at 10,000 g for 45-minutes (Optima LE-80 K ultracentrifuge, JA25.50 rotor, +4 °C, acceleration setting: max, deceleration setting: max) and then at 100,000 g for 90-minutes (Beckman Optima XE ultracentrifuge, Ti45 rotor, +4 °C, acceleration setting: max, deceleration setting: 7) and again for 90-minutes to wash the pellet with 0.22 μm filtered phosphate buffered saline (PBS, Sigma, D8662). As a negative control of EV collection, culture dishes without cells were processed in the same way.

For initial experiments, only ultracentrifugation isolation was used, but for the majority of experiments reported here, after removing the supernatant by pipetting, individual EV pellets were re-suspended in 100 μL of 0.22 μm filtered PBS, combined, and the total volume was adjusted to 300 μL. The concentrated sample was then loaded onto an IZON qEV single size-exclusion chromatography column (70 nm, 1001035), which had been prepared by removing the buffer from the top and washing the column twice with filtered PBS. Sample application was followed immediately by fraction collection. Once the sample passed through, two drops of filtered PBS were added to the column, followed by additional PBS to ensure flow. Fractions were collected as follows: the first void volume (800 μL, discarded), the EV-containing fraction (800 μL, retained), the second void (1200 μL, discarded), and the protein fraction (1800 μL, retained if needed). EV-containing fractions were stored at -80°C.

#### Nanoparticle tracking analysis (NTA)

EV characterization was performed according to the MISEV 2018 guidelines^33^. The particle concentration and size distribution from pellets and individual SEC fractions was measured using a NanoSight NS300 (Malvern, Worcestershire, UK). Each sample was diluted 1:50 or more with PBS that had been verified to be free of contaminants (i.e., no particles detected by NanoSight NS300) to achieve a final concentration between 5×10^7^ and 9×10^8^ particles/mL. Each 400 μL sample was injected into the sample chamber at a constant flow rate using the Malvern NanoSight syringe pump system. For all recordings, the camera level was adjusted to 15 and three 60 s videos were recorded for each sample. For analysis, the detection threshold was set to 5 and the other settings were “automatic”. For subsequent EV-related experiments, a working concentration of 10^6^ EVs/mL was used.

#### Transmission electron microscopy (TEM)

Copper-carbon film grids (400 mesh; EM Resolutions, Sheffield, United Kingdom) were glow-discharged using a Quorum K100X glow discharger. Grids were then placed on a 5 μL droplet of sample (on dental wax) for 2 min. Buffer salts were removed by transferring the grids twice to a fresh drop of distilled water and incubated for 5 s each. The excess fluid was removed with filter paper, and the grids were transferred to one drop of uranyl acetate (1.5% in distilled water) and incubated for 1-minute. Excess fluid was removed with filter paper and the grids were air dried prior to imaging. Grids were imaged using an FEI/ThermoFisher Scientific Tecnai G2 TEM electron microscope run at 200 keV accelerating voltage and using a 20 μm objective aperture to improve contrast. Images were acquired with an ORCA HR high resolution CCD camera using a Hamamatsu DCAM board and running the Image Capture Engine, software version 600.323, from AMT Corp. (Advanced Microscopy Techniques Corp. Danvers, USA).

#### Proteomic analysis of MSC-EVs

Enriched MSC-EV samples were incubated with 1 mL ice-cold acetone overnight at -20°C. Precipitated proteins were pelleted by centrifugation at 13,500 g (Beckman Optima XE ultracentrifuge, Ti45 rotor, +4 °C, acceleration setting: max, deceleration setting: 7). Pellets were resuspended in 50 μL NuPAGE LDS sample buffer (NP0007, Thermo Fisher Scientific) and boiled at 95 °C for 5-minutes. A 25 μL sample was loaded onto a precast NuPAGE 4% to 12% Bis-Tris polyacrylamide gel (Thermo Fisher Scientific) and electrophoresed for 1.5 cm at 75V. Following electrophoresis, the gel was washed in deionized water and stained with SimplyBlue SafeStain (LC6060, Thermo Fisher Scientific) to visualize total protein bands. Each lane was then excised and cut into two equal-sized pieces for in-gel digestion. Gel pieces were destained using 50% acetonitrile in 50 mM ammonium bicarbonate until clear. Proteins were reduced with 10 mM dithiothreitol in 50 mM ammonium bicarbonate at 56 °C for 45-minutes, followed by alkylation with 55 mM iodoacetamide in 50 mM ammonium bicarbonate at room temperature in the dark for 30-minutes. After additional washes and dehydration with acetonitrile, the gel pieces were rehydrated with 25 μg sequencing-grade modified trypsin (11418025001, Sigma) in 50 mM ammonium bicarbonate and incubated overnight at 37 °C for protein digestion. The resulting tryptic peptides were extracted, dried using a vacuum centrifuge and resuspended in mass spectrometry solvent (0.1% TFA, 3% MeCN) for liquid chromatography with tandem mass spectrometry (LC-MS/MS) analysis.

LC-MS/MS was performed on a Q Exactive Plus interfaced with an EASY-spray source coupled to an RSLC3000 nanoU-PLC (Thermo Fisher Scientific). Tryptic peptides were fractionated by a 50 cm C18 PepMap column (Thermo Fisher Scientific) maintained at 40 °C with solvent A (0.1% formic acid) and solvent B (80% MeCN, 0.1% formic acid), utilizing a gradient rising from 3% to 40% solvent B between 7- and 52-minutes followed by a wash at 95% B and 25-minutes column re-equilibration at 3% solvent A. A spray voltage of 1.3 kV was used with MS spectra acquired from 400 to 1500 m/z at 70,000 resolution. MS/MS spectra were acquired in a top 10 fashion at 17,500 resolution with a target ACG of 1×10^5^ and a maximum injection time of 250 milliseconds. Data were processed using MaxQuant v.2.0.1.0 with *Homo sapiens* database (Proteome ID: UP000005640) with label-free quantitation and iBAQ enabled.

#### EV surface protein shaving

Two methods were used to remove protein from the external EV surface, so-called shaving, to perturb EV internalization and surface membrane receptor-ligand interactions:

##### Trypsin

Trypsin hydrolysis of EV surface proteins was performed as previously described^82^. Briefly, trypsin solution was prepared by dissolving 25 μg of sequencing-grade modified trypsin (Roche 11418025001) in 250 μL of 50 mM ammonium bicarbonate (prepared by dissolving 0.395 g ammonium bicarbonate in 100 mL ddH_2_O), resulting in a final concentration of 0.1 μg/μL. An equal volume of the trypsin solution was added to EV samples (specifically the UC-SEC fraction 3), and the mixture was incubated at 37°C in a hot block for 5-minutes. For ultracentrifugation (UC) samples, an additional incubation of 30-minutes was performed. Samples were then transferred to an ultracentrifugation rotor and centrifuged at 100,000 × g (Beckman Optima XE ultracentrifuge, Ti45 rotor, +4 °C, acceleration setting: max, deceleration setting: 7) for 1-hour at 4°C. Following centrifugation, the EV pellet was resuspended in 50 μL of sterile PBS.

##### Proteinase K

Proteinase K digestion of EV surface proteins was performed as previously described^41^. EV-PBS solution was diluted in 1 mL of sterile PBS. The appropriate volume being determined by the previous NTA result. For proteinase K treatment, 750 μL of UC-SEC-derived EVs was incubated with proteinase K at a final concentration of 25 μg/mL for 1-hour at 37°C, while the remaining 250 μL was reserved as an untreated control. The enzymatic reaction was stopped by adding 1 μL of 250 mM phenylmethylsulfonyl fluoride (PMSF) prepared in DMSO. Samples were then transferred to an ultracentrifugation rotor and centrifuged at 100,000 × g (Beckman Optima XE ultracentrifuge, Ti45 rotor, +4 °C, acceleration setting: max, deceleration setting: 7) for 1-hour at 4°C. The resulting EV pellet was resuspended in 50 μL of sterile PBS.

#### PKH26 staining of EVs

All EV samples (trypsin-treated EVs, proteinase K-treated EVs, and untreated control EVs) were thawed on ice and briefly vortexed before being diluted in PBS to a final volume of 250 μL. EVs were then labelled using the PKH26 Red Fluorescent Cell Linker Kit (Sigma-Aldrich, PKH26GL), following the manufacturer’s instructions. Briefly, 1 μL of PKH26 dye was added to 250 μL of Diluent C to create the working dye solution, which was immediately mixed with 250 μL EV solution by pipetting. The mixture was incubated at room temperature for 5-minutes with periodic mixing. To quench excess dye, 500 μL of 1% BSA was added and incubated for 1-minute. The labelled EVs were then transferred to an ultracentrifuge tube and topped up with 63 mL of 1% BSA. Ultracentrifuge tubes were balanced and centrifuged at 100,000 × g for 90-minutes at 4°C. The final EV pellet was resuspended in 100 μL of PBS for downstream NTA to determine the size distribution and concentration.

#### Confocal microscopy

DRG neurons were cultured overnight prior to the experiment. Before confocal imaging, PKH26-labelled EVs were added to the DRG neuron cultures at a final concentration of 10^6^ EVs/mL in 2 mL of culture medium (concentration determined by NTA), and neurons incubated with EVs at 37 °C for 10-minutes.

Confocal microscopy was performed using a Zeiss LSM 980 laser scanning microscope (Carl Zeiss AG, Germany) equipped with an Airyscan 2 detection unit. Imaging was conducted with a Plan-Apochromat 63×/1.40 NA oil DIC M27 objective (Zeiss) using Immersol 518 F immersion oil (refractive index n = 1.518 at 23 °C). Image acquisition and processing were carried out using ZEN (Zeiss) and Zeiss Arivis Pro software. Fluorescence from PKH26-labelled EVs was imaged using both conventional confocal and Airyscan super-resolution modes. Images were acquired at 12-bit depth with a resolution of 1024 × 1024 pixels (pixel size = 0.02 μm), a zoom of 3, sequential and unidirectional scanning, line averaging (n = 2), and a pinhole size of 1 AU for confocal mode. Excitation was performed using a 639 nm laser at 2% intensity, and emission was collected between 540–600 nm. Detector gain and dwell time (gain = 729; dwell time = 12.6 μs) were optimized to minimize signal saturation and photobleaching. Z-stack images were acquired using a step size of 0.6 μm, automatically calculated by Zeiss Arivis Pro software to meet Nyquist sampling criteria, resulting in 50-120 optical slices per stack. Maximum intensity projections and 3D reconstructions were generated using Zeiss Arivis Pro software. For Airyscan data, the raw signal from the 32-detector array was processed via filtering, deconvolution, and pixel reassignment to enhance spatial resolution and signal-to-noise ratio.

#### Machine learning model and supervised segmentation

Segmentation of EVs and DRG neurons from image stacks was performed using the machine learning-based trainable segmentation plugin within the Zeiss Arivis Pro software. Prior to training, images were pre-processed using edge detection and noise reduction techniques, including Gaussian blur and membrane projection filters, to enhance structural boundaries. EVs, identified by PKH26 fluorescence, were manually annotated to distinguish them from background signals.

For model training, approximately 200, 100 and 100 elements were manually classified within each region of interest (ROI, corresponding to DRG neurons), representing three categories: “EVs,” “background,” and “gaps” (the peripheral, partially faded regions demarcating EVs from background). Classifiers were trained for each image stack using these features within an ROI. The resulting classifiers were then applied to their respective ROIs to generate multi-channel probability maps.

All generated probability maps were projected into 3D using Zeiss Arivis Pro, with each segmentation class assigned to a distinct image channel. The EV channel from the final probability maps was selected for 3D visualization and further analysis.

#### Image analysis

The number of internalized EVs and the volume of DRG neurons were quantified. To assess whether the surface-shaving protocol reduced EV uptake, the average number of EVs per 1 μm^3^ of neuronal volume was calculated. A one-way ANOVA followed by Tukey’s post hoc test was used to compare the three experimental groups (i.e. trypsin-treated, proteinase K-treated and control EVs). Statistical significance was defined as *p* ≤ 0.05. Data are presented as mean ± SEM.

### MSC-EV: miR assay

#### RNA extraction, real-time PCR and small RNA sequencing

RNA from purified MSC-EVs was extracted using TRIzol, followed by using the RNA Clean and concentrator kit (Zymo Research, # R1013). Briefly, TRIzol-sample mixes were incubated for 30-minutes, then they were combined with 0.2 volume of chloroform:isoamyl, and left for 5-minutes at room temperature. Solutions were centrifuged (12,000 × *g*, 15-minutes, 5 °C); the aqueous phase was collected and combined the first buffer of the RNA Clean and concentrator kit and then follow the protocol according manufacturer’s instructions. Finally, the total RNA was eluted in 12x μL of nuclease-free water.

For small RNA sequencing, RNA integrity was measured using an Agilent 2100 Bioanalyzer Small RNA kit (5067-1548, Agilent Technologies, USA) for total RNA (RNA nano-chips) and for small RNA (small RNA chips), concentrations were measured in a Qubit 2.0 Fluorometer and using the Qubit RNA HS assay kit). Small RNA libraries were prepared and sequenced by Novogene. Human exosome small RNA-seq (WOBI) was performed, generating 20 million reads per sample using Illumina single-end 50 bp (SE50) sequencing.

#### Bioinformatics analyses

Bioinformatic analyses were performed with R version 4.2.1. The raw reads including the middle linker sequence were filtered from fastq files using the command line, and the number of unique molecular identifiers (UMIs) for sequences longer than 15 nucleotides (nt) in each sample were calculated using a self-written R-script. The results from the two obtained libraries were compared with PCA and correlation analyses and as a result, the count numbers of both libraries (“miRNA” and “long”) were combined for individual samples.

For the analysis of mature miRNAs, the data were further filtered to contain only 16-24 nt read sequences matching mature miRNAs. Read alignment and differential expression analysis were performed using scripts from DEUS 1.0 pipeline. At first, the remaining 16-24 nt long sequences were aligned to mature human miRNAs from miRbase release 22.1 using blast-based alignment allowing one mismatch and the count numbers for sequences aligned to the same miRNA were combined. Not-aligned sequences were discarded. Differential expression analysis was performed separately for EV and protein fractions, combining HC and UTI groups as a control group, and comparing with the UC samples. During the analysis based on DEseq2 (Version 1.36.0), replicate samples (libraries prepared from the same RNA) were collapsed together.

#### Gold nanoparticle functionalization and DRG neuron delivery

To mimic the effect of a miRNA delivery from EVs, we conjugated gold nanoparticles with RNA duplexes corresponding to each candidate miRNA. Citrate-stabilized gold nanoparticles with a diameter of 10 nm (Sigma, 741957) were prepared in double-distilled water (ddH₂O). Each duplex consisted of a guide strand, representing the mature miRNA sequence, and a complementary passenger strand. Since the passenger strand is typically discarded during RISC loading, the strand was modified with a 3′-terminal thiol linker to facilitate covalent attachment to the gold nanoparticle surface. No fluorescent tag was incorporated in this construct.

Four gold nanoparticle-miRNA duplex conditions were tested. The first carried miR-21-5p (miR-21-5p-duplex-THIOL), composed of a guide strand (5′-UAGCUUAUCAGACUGAUGUUGA-3′) and a passenger strand (5′-UCAACAUCAGUCUGAUAAGCUA[dT]_5_-thiol modifier C3 S-S-3′). The second carried miR-148-3p (miR-148-3p-duplex-THIOL), composed of a guide strand (5′-UCAGUGCACUACAGAACUUUGU-3′) and a passenger strand (5′-UCAACAUCAGUCUGAUAAGCUA[dT]_5_-thiol modifier C3 S-S-3′). The third carried miR-451a (miR-451a-duplex-THIOL), composed of a guide strand (5′-AAACCGUUACCAUUACUGAGUU-3′) and a passenger strand (5′-AACUCAGUAAUGGUAACGGUUU-[dT]₅-thiol modifier C3 S–S–3′). The fourth condition served as a negative control and was functionalized with a non-targeting sequence (Cy5NP) used to assess nanoparticle uptake by DRG neurons. This control duplex contained a Cy5 fluorophore at the 5′ end and a thiol-modified 3′ terminus (/Cyanine5/TTTTTTTTTTTTTTTTTTTTTTTTTTTTTTTTTTTTTTTT/5ThioC6/), enabling fluorescent tracking, but lacking any miRNA-like activity.

For gold nanoparticle functionalization, 3 mL of 1 OD gold nanoparticles was incubated with modified oligonucleotides (5 pmol/μL) and Tris(2-carboxyethyl) phosphine (TCEP, 100 equivalents per oligo) for 2h RT. Then, 12 μL of a 5 M NaCl solution were added every 20 min (5 times) until 0.3 M salt concentration was reached in the solution. The mixture was incubated for 16 h in the dark at room temperature. The gold nanoparticles were then centrifuged at 800,000 rpm at 4 °C for 45-minutes, the supernatant was removed, and the pellet was resuspended in water. This process was repeated twice to remove the unbound oligonucleotides.

To quantify the attached oligonucleotide, the supernatants were collected, concentrated, and quantified by spectroscopy at 260 nm using a standard curve. Gold nanoparticles functionalized with miRNAs or Cy5 control were prepared at final concentrations of 3.15 μM (miR-21-5p), 2.03 μM (miR-148-3p), 1.97 μM (miR-451a), 3.47 μM (MiRmix), and 0.41 μM (Cy5 control), respectively. 1 nM AuNPs were mixed with pre-warmed DRG culture medium and applied directly to the neuronal cultures for treatment.

## Supplementary information

**Supplementary Fig. 1:**
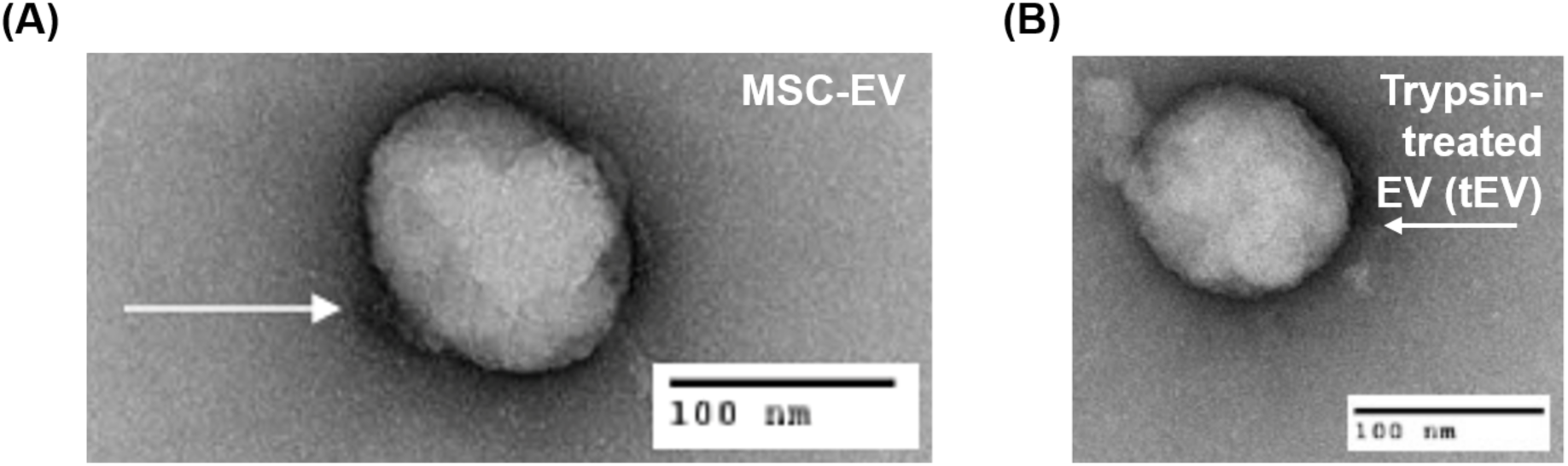
Transmission electron microscopy (TEM) images of (**a**) MSC-EV, (**b**) trypsin-treated EV (tEV) show that enzymatic shaving has minimal impact on EV membrane integrity.

**Supplementary Table 1.**
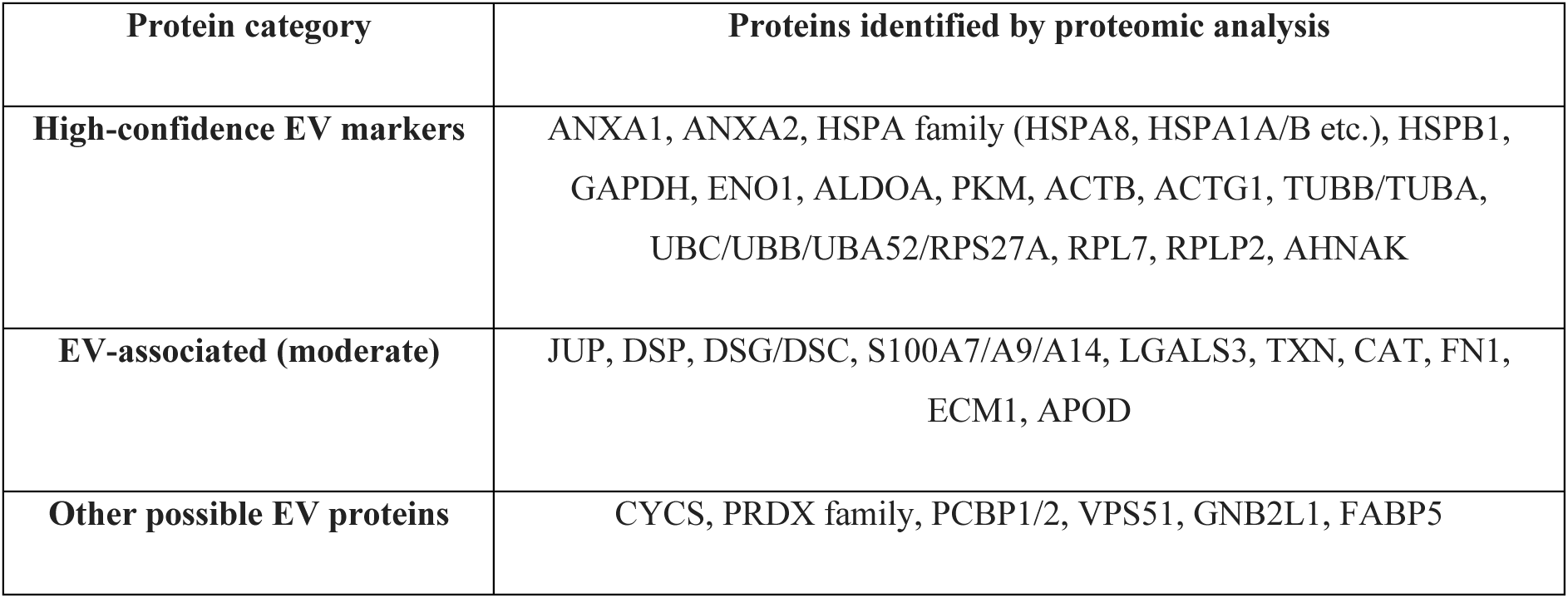
Summary of representative extracellular vesicle marker proteins identified by proteomic analysis.

**Supplementary Table 2.** Effects of MSC-EVs on intrinsic and active properties of NGF-sensitized IB4-ve DRG neurons.

|  | Vehicle<br>(n=16) | NGF<br>(n=20) | NGF+EV<br>(n=14) |
| --- | --- | --- | --- |
| Capacitance (pF) | 19.12±1.04 | 17.98±0.76 | 16.24±1.12 |
| Resting membrane potential (RMP, mV) | -53.90±1.60 | -40.51±2.45 * | -51.18±0.87 † |
| Rheobase (pA) | 551.88±45.94 | 228.50±30.87 * | 485.71±74.55 † |
| Peak amplitude (mV) | 48.98±2.43 | 51.88±1.61 | 52.39±2.62 |
| Half-peak duration (ms) | 4.63±0.64 | 8.06±0.53 * | 4.91±0.81 † |
| Afterhyperpolarization (AHP) amplitude (mV) | 11.44±1.69 | 10.73±1.63 | 11.08±1.87 |
| AHP duration (AHP tau, ms) | 6.32±0.79 | 10.25±0.88 * | 8.69±0.95 |

Abbreviations: NGF – nerve growth factor; EV – extracellular vesicle. Data were analyzed using ordinary one-way ANOVA with Bonferroni post-hoc analysis. Data are displayed as mean±SEM. \* p-adj < 0.05 compared to vehicle group. † p-adj < 0.05 compared to NGF group. For clarity of reading, significance is denoted when p < 0.05, rather than the actual significance level being given for each comparison.

**Supplementary Table 3.** Particle distribution of trypsin-treated MSC-EVs (tEVs) and Proteinase K-treated MSC-EVs (pkEVs)

|  | Untreated MSC-EVs | Trypsin-treated MSC-EVs (tEVs) | Proteinase K-treated MSC-EVs (pkEVs) |
| --- | --- | --- | --- |
| Size range (nm) | 80-520 | 50-420 | 20-320 |
| Peak(s) (nm) | 152 | 100 and 162 | 129 |
| Mean size (nm) | 176.8 | 122.8 | 142.3 |
| Standard deviation (nm) | 71.1 | 51.9 | 45.5 |

**Supplementary Table 4.** Effects of trypsin-treated MSC-EVs (tEVs) on intrinsic and active properties of NGF-sensitized IB4-ve DRG neurons.

|  | Vehicle<br>(n=16) | NGF<br>(n=19) | NGF+EV<br>(n=17) | NGF+tEV<br>(n=22) |
| --- | --- | --- | --- | --- |
| Capacitance (pF) | 15.69±0.91 | 17.38±0.72 | 16.38±0.78 | 16.99±0.53 |
| Resting membrane potential (RMP, mV) | -57.18±0.99 | -39.38±2.92 * | -55.67±0.71 † | -49.12±1.36 * † § |
| Rheobase (pA) | 450.63±50.27 | 248.42±32.98 * | 452.84±38.05 † | 300.00±30.72 * § |
| Peak amplitude (mV) | 55.66±2.24 | 49.54±1.28 | 51.03±2.50 | 53.84±1.42 |
| Half-peak duration (ms) | 4.50±0.21 | 7.55±0.70 * | 5.91±0.97 | 4.58±0.37 † |
| Afterhyperpolarization (AHP) amplitude (mV) | 7.19±1.38 | 11.74±1.52 | 8.86±1.16 | 12.08±1.21 |
| AHP duration (AHP tau, ms) | 5.45±0.56 | 9.36±0.94 * | 7.67±0.95 | 9.19±0.69 * |

Abbreviations: NGF – nerve growth factor; EV – extracellular vesicle. Data were analyzed using ordinary one-way ANOVA with Bonferroni post-hoc analysis. Data are displayed as mean±SEM. \* p-adj < 0.05 compared to vehicle group. † p-adj < 0.05 compared to NGF group. § p-adj < 0.05 compared to NGF+EV group. For clarity of reading significance is denoted when p < 0.05, rather than the actual significance level being given for each comparison.

**Supplementary Table 5.** Effects of proteinase K-treated MSC-EVs (pkEVs) on intrinsic and active properties of NGF-sensitized IB4-ve DRG neurons.

|  | Vehicle<br>(n=15) | NGF<br>(n=10) | NGF+EV<br>(n=7) | NGF+pkEV<br>(n=18) |
| --- | --- | --- | --- | --- |
| Capacitance (pF) | 16.04±0.90 | 15.48±1.09 | 15.72±0.77 | 15.60±0.75 |
| Resting membrane potential (RMP, mV) | -56.60±1.18 | -40.20±2.65 * | -56.75±0.90 † | -48.90±0.93 * † § |
| Rheobase (pA) | 569.33±59.20 | 256.00±30.56 * | 531.43±65.08 † | 317.78±22.56 * § |
| Peak amplitude (mV) | 48.29±1.71 | 48.04±3.21 | 50.01±3.03 | 49.83±1.32 |
| Half-peak duration (ms) | 7.79±0.48 | 6.99±1.16 | 5.64±1.27 | 6.41±0.56 |
| Afterhyperpolarization (AHP) amplitude (mV) | 10.01±0.88 | 14.79±2.33 | 8.76±2.08 | 12.37±0.93 |
| AHP duration (AHP tau, ms) | 5.91±0.46 | 10.59±1.62 * | 7.77±2.08 | 8.41±0.88 |

Abbreviations: NGF – nerve growth factor; EV – extracellular vesicle. Data were analyzed using ordinary one-way ANOVA with Bonferroni post-hoc analysis. Data are displayed as mean±SEM. \* p-adj < 0.05 compared to vehicle group. † p-adj < 0.05 compared to NGF group. § p-adj < 0.05 compared to NGF+EV group. For clarity of reading significance is denoted when p < 0.05, rather than the actual significance level being given for each comparison.

**Supplementary Table 6.** Effects of 10-minute treatment of MSC-EVs on intrinsic and active properties of NGF-sensitized IB4-ve DRG neurons.

|  | Vehicle<br>(n=20) | NGF<br>(n=19) | NGF+10-min EV<br>(n=21) |
| --- | --- | --- | --- |
| Capacitance (pF) | 15.94±0.63 | 15.20±0.70 | 14.17±0.72 |
| Resting membrane potential<br>(RMP, mV) | -51.97±1.05 | -45.80±1.44 * | -47.14±1.52 * |
| Rheobase (pA) | 441.05±45.94 | 264.50±37.70 * | 299.05±40.66 * |
| Peak amplitude (mV) | 50.07±1.27 | 50.39±1.27 | 52.82±1.93 |
| Half-peak duration (ms) | 5.14±0.77 | 8.52±1.03 * | 7.86±0.68 |
| Afterhyperpolarization<br>(AHP) amplitude (mV) | 8.56±0.99 | 13.07±1.29 * | 10.50±1.23 |
| AHP duration (AHP tau, ms) | 7.02±0.75 | 10.60±0.89 * | 8.08±0.65 |

Abbreviations: NGF – nerve growth factor; EV – extracellular vesicle. Data were analyzed using ordinary one-way ANOVA with Bonferroni post-hoc analysis. Data are displayed as mean±SEM. For clarity of reading significance is denoted as \* when $p < 0.05$ , rather than the actual significance level being given for each comparison.

**Supplementary Table 7.**
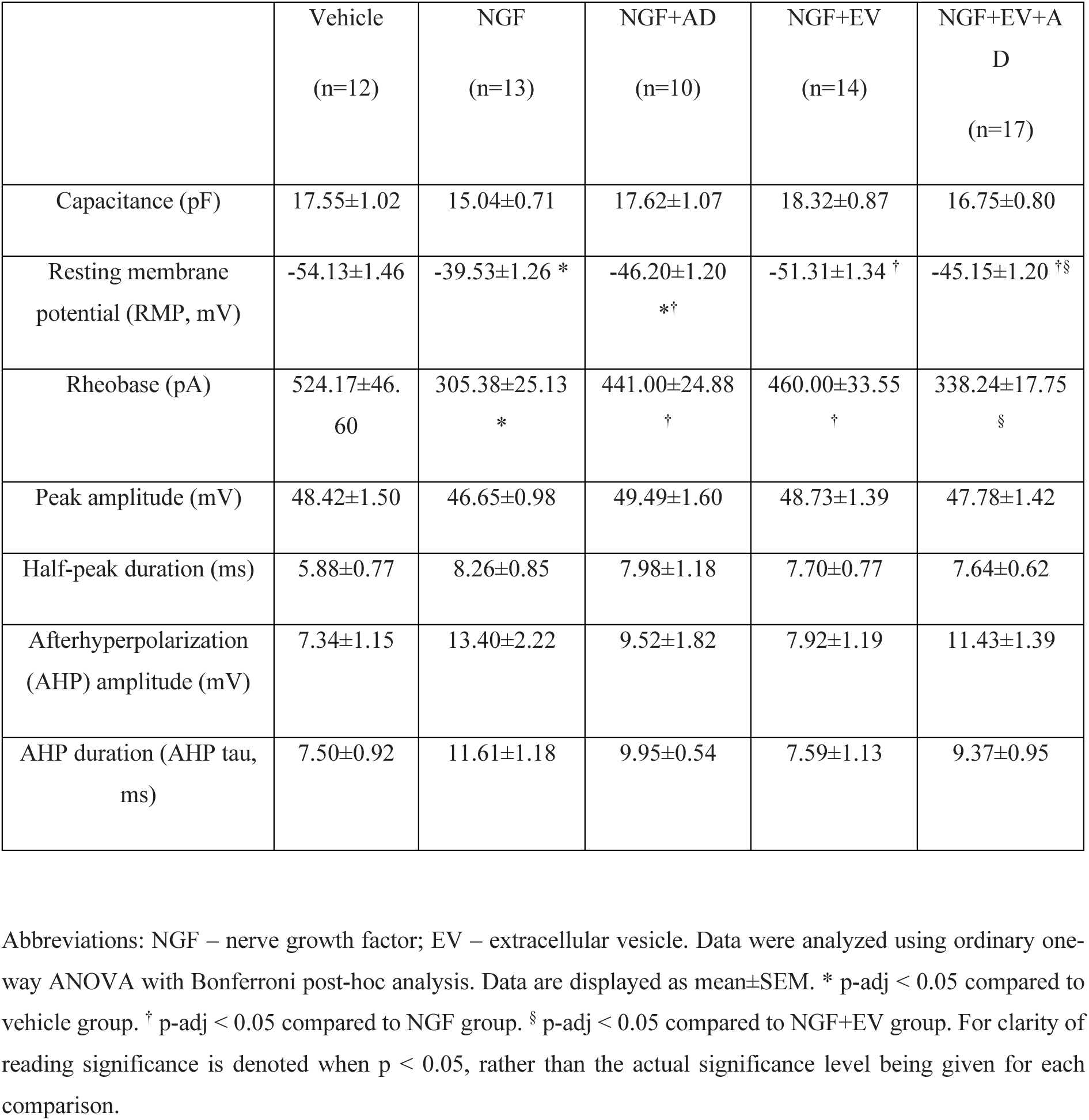
Effects of actinomycin D of MSC-EVs on intrinsic and active properties of NGF-sensitized IB4-ve DRG neurons.

**Supplementary Table 8.**
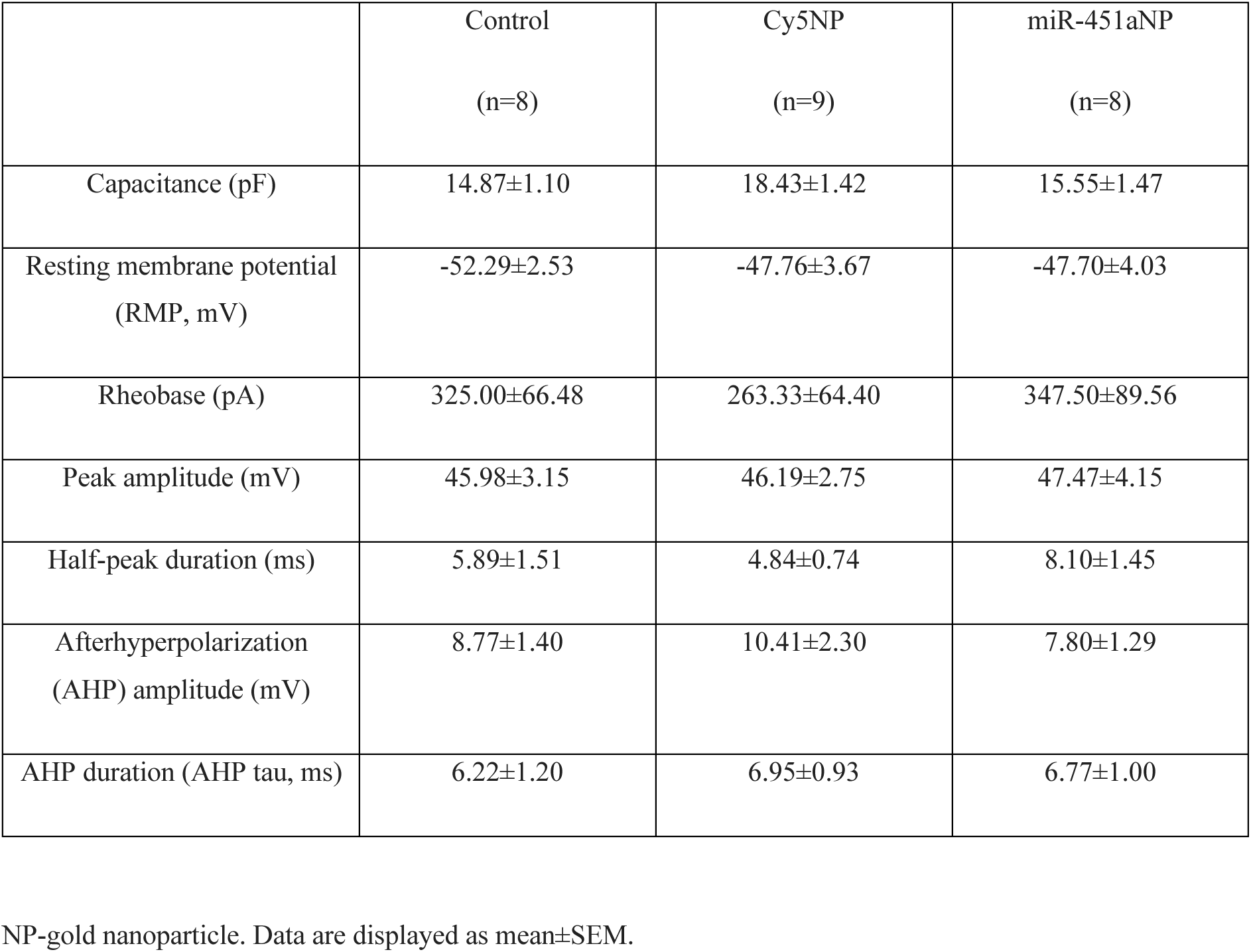
Effects of Cy5- and miR-451a-conjugated gold nanoparticles on intrinsic and active properties of on intrinsic and active properties of DRG neurons.

**Supplementary Table 9.**
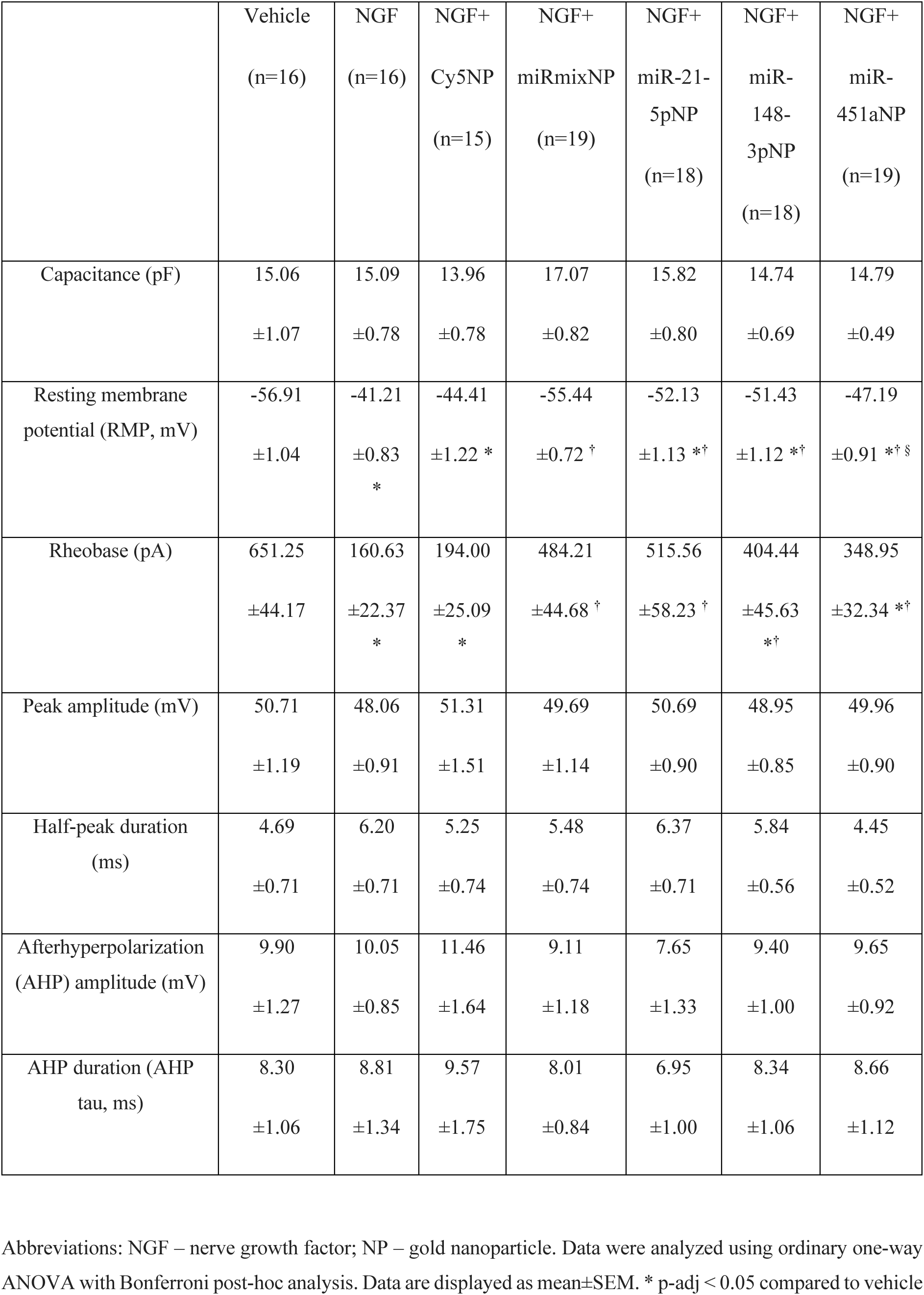
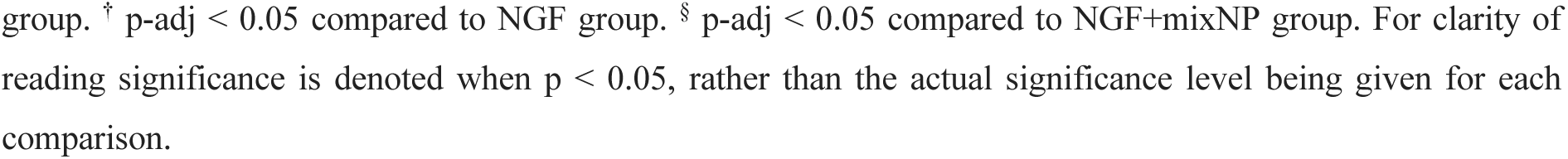
Effects of miR-conjugated gold nanoparticles on intrinsic and active properties of NGF-sensitized IB4-ve DRG neurons.

|  | Vehicle<br>(n=16) | NGF<br>(n=16) | NGF+<br>Cy5NP<br>(n=15) | NGF+<br>miRmixNP<br>(n=19) | NGF+<br>miR-21-<br>5pNP<br>(n=18) | NGF+<br>miR-<br>148-<br>3pNP<br>(n=18) | NGF+<br>miR-<br>451aNP<br>(n=19) |
| --- | --- | --- | --- | --- | --- | --- | --- |
| Capacitance (pF) | 15.06<br>±1.07 | 15.09<br>±0.78 | 13.96<br>±0.78 | 17.07<br>±0.82 | 15.82<br>±0.80 | 14.74<br>±0.69 | 14.79<br>±0.49 |
| Resting membrane<br>potential (RMP, mV) | -56.91<br>±1.04 | -41.21<br>±0.83<br>* | -44.41<br>±1.22 * | -55.44<br>±0.72 † | -52.13<br>±1.13 *† | -51.43<br>±1.12 *† | -47.19<br>±0.91 *†§ |
| Rheobase (pA) | 651.25<br>±44.17 | 160.63<br>±22.37<br>* | 194.00<br>±25.09<br>* | 484.21<br>±44.68 † | 515.56<br>±58.23 † | 404.44<br>±45.63<br>*† | 348.95<br>±32.34 *† |
| Peak amplitude (mV) | 50.71<br>±1.19 | 48.06<br>±0.91 | 51.31<br>±1.51 | 49.69<br>±1.14 | 50.69<br>±0.90 | 48.95<br>±0.85 | 49.96<br>±0.90 |
| Half-peak duration<br>(ms) | 4.69<br>±0.71 | 6.20<br>±0.71 | 5.25<br>±0.74 | 5.48<br>±0.74 | 6.37<br>±0.71 | 5.84<br>±0.56 | 4.45<br>±0.52 |
| Afterhyperpolarization<br>(AHP) amplitude (mV) | 9.90<br>±1.27 | 10.05<br>±0.85 | 11.46<br>±1.64 | 9.11<br>±1.18 | 7.65<br>±1.33 | 9.40<br>±1.00 | 9.65<br>±0.92 |
| AHP duration (AHP<br>tau, ms) | 8.30<br>±1.06 | 8.81<br>±1.34 | 9.57<br>±1.75 | 8.01<br>±0.84 | 6.95<br>±1.00 | 8.34<br>±1.06 | 8.66<br>±1.12 |

Abbreviations: NGF – nerve growth factor; NP – gold nanoparticle. Data were analyzed using ordinary one-way ANOVA with Bonferroni post-hoc analysis. Data are displayed as mean±SEM. \* p-adj < 0.05 compared to vehicle
group. <sup>†</sup> p-adj < 0.05 compared to NGF group. <sup>§</sup> p-adj < 0.05 compared to NGF+mixNP group. For clarity of reading significance is denoted when $p < 0.05$ , rather than the actual significance level being given for each comparison.

## Notes

### Competing Interest Statement

The authors have declared no competing interest.

